# Integrative Modeling of Protein-Polypeptide Complexes by Bayesian Model Selection using AlphaFold and NMR Chemical Shift Perturbation Data

**DOI:** 10.1101/2024.09.19.613999

**Authors:** Tiburon L. Benavides, Gaetano T. Montelione

## Abstract

Protein-polypeptide interactions, including those involving intrinsically-disordered peptides and intrinsically-disordered regions of protein binding partners, are crucial for many biological functions. However, experimental structure determination of protein-peptide complexes can be challenging. Computational methods, while promising, generally require experimental data for validation and refinement. Here we present *CSP_Rank*, an integrated modeling approach to determine the structures of protein-peptide complexes. This method combines AlphaFold2 (AF2) enhanced sampling methods with a Bayesian conformational selection process based on experimental Nuclear Magnetic Resonance (NMR) Chemical Shift Perturbation (CSP) data and AF2 confidence metrics. Using a curated dataset of 108 protein-peptide complexes from the Biological Magnetic Resonance Data Bank (BMRB), we observe that while AF2 typically yields models with excellent consistency with experimental CSP data, applying enhanced sampling followed by data-guided conformational selection routinely results in ensembles of structures with improved agreement with NMR observables. For two systems, we cross-validate the CSP-selected models using independently acquired nuclear Overhauser effect (NOE) NMR data and demonstrate how CSP and NMR can be combined using our Bayesian framework for model selection. *CSP_Rank* is a novel method for integrative modeling of protein-peptide complexes and has broad implications for studies of protein-peptide interactions and aiding in understanding their biological functions.

## INTRODUCTION

Structural analysis of protein-peptide complexes is pivotal for understanding biological mechanisms and facilitating development of peptide and peptide-mimicking therapeutics (Sugase et al, 2007; Watkins & Arora, 2015; Lee et al, 2019). In favorable cases, structures of protein-peptide complexes can be determined using X-ray crystallography, cryogenic electron microscopy (cryoEM), or Nuclear Magnetic Resonance spectroscopy (NMR). NMR offers several advantages, including the ability to examine the molecule of interest in a near-native solution environment and to provide quantitative information about conformational dynamics. However, experimental structure determination of protein-peptide complexes by conventional NMR methods involves substantial effort and generally requires isotope-enrichment of both the protein receptor and the polypeptide ligand (Aiyer et al, 2021).

Recent advances in computational structure prediction methods now frequently deliver high-quality protein-peptide docking poses with accuracy often comparable with experimental studies (Johansson-Åkhe & Wallner, 2022; Tsaban et al, 2022; Lensink et al, 2023, Ozden et al, 2023;). Despite the longstanding challenge of predicting optimal pairwise protein docking poses (Janin et al, 2003), recently developed atomic-resolution protein structure prediction tools such as *AlphaFold2-multimer* (AF2) (Jumper et al, 2021; Evans et al, 2021), *AlphaFold3* (AF3) (Abramson et al, 2024), *RosettaFold* (Baek et al, 2021; Krishna et al, 2024) and *Evolutionary Scale Modeling* (ESMFold) (Lin et al, 2023) have reinvigorated the field. These tools frequently outperform more traditional protein-peptide docking algorithms especially in cases involving extensive structural changes, such as those that occur upon binding of disordered polypeptides to receptor proteins (Ko & Lee, 2021; Tsaban et al, 2022; Zhang et al, 2023; Mondal et al, 2023). Although the evolving machine learning (ML)-based modeling methods often provide remarkably accurate protein structure predictions (Jumper et al, 2021; Huang et al, 2021; Bryant et al, 2022; Tejero et al, 2022; Li et al, 2023), in some cases AF2 structure models do not match to experimentally-determined structures (Huang et al, 2021; Terwilliger et al, 2022; Bonin et al, 2024; Huang & Montelione, 2024). These counter examples provide important lessons about the strengths and weaknesses of ML-based modeling methods and demonstrate the need for experimental validation or refinement of ML-based models, particularly for flexible proteins and those that adopt multiple conformational states.

“Enhanced Sampling” (ES) describes a suite of emerging methods that modulate the data pipeline of ML-based structure prediction methods like AF2 to explore the conformational variability of protein structures. These methods are grounded in the hypothesis that either AF2 has learned by some kind of memorization the alternative conformations of protein structures (Del Alamo et al, 2022; Sala et al, 2023, Chakravarty et al, 2024), or the stronger claim that AF2 has learned insights into the free energy landscape governing protein conformational diversity (Roney & Ovchinnikov, 2022; Monteiro da Silva, 2023; Vani et al, 2023; Feng et al, 2024, Bryant & Noé, 2024). ES methods can be leveraged to predict conformational dynamics and alternative conformational states of biomolecules and possibly provide information about the pathways between disparate states (Kalakoti & Wallner, 2024).

Recent studies underscore the efficacy of ES techniques, including excellent performance in prediction of protein complexes (Lensink et al, 2023; Ozden et al, 2023; Wallner, 2023a; Wallner, 2023b) and in modeling alternative conformational states of proteins (Del Alamo et al, 2022; Stein & McHaourab, 2022; Kryshtafovych et al, 2023; Bryant & Noé, 2024; Chakravarty et al, 2024; Kalakoti & Wallner, 2024; Monteiro da Silva et al, 2024; Stein & McHaourab, 2024; Sala et al, 2023; Wayment-Steele et al, 2024). These methods rely on perturbing the input multiple sequence alignment (MSA) and/or varying weights of neural network nodes to achieve broader conformational sampling. *AFAlt* (Del Alamo et al, 2022) and *AFCluster* (Wayment-Steele et al, 2024) both use shallow MSAs to suppress dominance of evolutionary covariation information (ECs), which results in increased conformational diversity, allowing the modeling of alternative conformational states, while *SPEACH_AF* (Stein & McHaourab, 2024) suppresses ECs by MSA column masking. *AFSample* exploits alternative network weights and node dropouts to generate conformational diversity of AF2-predicted models (Wallner, 2023a), while *AFSample2* manipulates the MSA by performing alanine column masking like *SPEACH_AF*, but in a randomized manner that has been proposed to more robustly predict the transitional structures between two or more alternative states (Kalakoti & Wallner, 2024). ES methods can provide many biophysically plausible ‘shots on goal’ (i.e., varied and physically relevant structure models), improving the performance of AF2 in modeling complexes and in sampling conformational landscapes. ES methods can be used to generate collections of alternative conformations, providing the basis for *conformer selection* by ranking based on model consistency with experimental data (Huang & Montelione, 2024).

NMR chemical shift perturbations (CSPs) are changes in NMR resonance frequencies of atoms in a receptor protein due to binding by a ligand or biomolecular partner. They monitor both direct contacts between the receptor and ligand and conformational changes of the receptor, including long-range allosteric effects, due to ligand binding (Wishart, 2011; Williamson, 2013). They are highly informative and easily acquired for small, well-behaved proteins once backbone NMR resonance assignments have been determined. CSPs are routinely used in guiding NMR-based structural modeling of protein-protein and protein - nucleic acid complexes (Karaca & Bonvin, 2013). CSP-guided docking methods like *HADDOCK* often provide accurate structures of protein complexes using CSP data to guide the docking process (Dominguez et al, 2003; De Vries et al, 2010), particularly when CSPs are available for both partners in the interaction. *ColabDock*, a framework incorporating sparse distance restraints to guide AF2 modeling, has also been applied successfully in a few examples of CSP-based docking (Feng et al, 2024). However, interpreting CSPs as structural restraints presents challenges, particularly for docking of peptides which involve folding-upon-binding, and/or when ligand binding results in allosteric effects that perturb chemical shifts distant from the interaction interface (Schmitz et al, 2012; Nussinov & Tsai, 2015; Mondal et al, 2023; Skeens & Lisi, 2023). Additionally, CSPs encode information that may be averaged from multiple conformational states or complex assemblies, including multiple transient weak complexes which may have their own CSPs, complicating their interpretation as docking restraints (Schmitz et al, 2012; Robustelli et al, 2012; Mondal et al, 2023). Hence, significant challenges in integrative modeling using CSPs remain, especially for systems where CSPs can only be measured for one binding partner and those involving significant structural rearrangements of the receptor upon binding, where there are significant numbers of CSPs distant from the binding interface, or folding-upon-binding of the polypeptide ligand.

In this paper we present *CSP_Rank*, an integrative modeling protocol for determining the structures of protein - peptide complexes that combines multiple ES approaches like *AFSample, AFSample2*, and *AFAlt* with ligand-induced CSP data for the receptor protein. Using a database of 108 protein-peptide complexes with known 3D structures and available NMR chemical shift data, we observed that baseline *AF2-multimer* models of these complexes have excellent knowledge-based structure-quality features and achieve high structural similarity to the corresponding complex models deposited by experimental research groups in the Protein Data Bank (PDB). Using a metric of model quality based on comparing predicted and observed CSPs due to complex formation, many (but not all) AF2 models of these protein-peptide complexes score better than the corresponding experimental models available in the PDB. Using some of these protein-peptide complexes as test cases, we demonstrate the *CSP_Rank* protocol, which uses AF2 ES methods to generate collections of protein-peptide complex structures and ranks the resulting models using a Bayesian conformer selection score assessing the fit of models to CSP data and AF2 model reliability scores. The resulting ensembles of protein - peptide complexes usually fit experimental CSP data better than the corresponding structures available in the PDB or the baseline AF2-multimer models. In several cases the accuracy of the resulting *CSP_Rank* models was also cross-validated with experimental NMR NOESY data and were found to fit these data as well as structures generated using the NOESY data as restraints. Finally, we demonstrate how CSP and NOESY data can be combined using our Bayesian framework for model selection from the enhanced-sampled AF2 conformer distributions.

## METHODS

### Data collection

Chemical shift lists for holo(bound) and apo(free) forms of proteins were collected using the BioMagResBank (BMRB) API (Hoch et al, 2023). The criteria for inclusion in this set were binary complexes between an ordered protein receptor and a polypeptide of 80 amino-acid residues or less, with backbone resonance assignments available for both the apo and holo versions of the receptor protein. As additional criteria, the experimental conditions used in determining NMR assignments for the apo and holo protein receptor must be approximately the same in terms of ionic strength, pH (± 0.5 pH units), and temperature (± 5 degrees C). The representative structure of the bound form of the complex used for comparison was selected as the medoid model of the NMR ensemble deposited in the PDB, as determined with the program *PDBStat* (Tejero et al, 2013).

### Structure computation

Structures of both the free (apo) protein receptor and the bound (holo) protein-polypeptide complex were modeled using *AlphaFold Multimer v2*.*3* available on Google Colab (Mirdita et al, 2022) using no templates and three recycles (hereafter denoted as ‘baseline AF2’). Only the first-ranked model of five models returned by baseline AF2 were used for comparison with the medoid PDB model. AF2 models were protonated using the Amber relax Google Colab notebook protocol (Mirdita et al, 2022).

For several protein - polypeptide complexes, we also computed protein-peptide complex structures using the ES methods *AFSample* (Johansson-Åkhe & Wallner, 2022), *AFSample2* (Kalakoti & Wallner, 2024), and/or *AFAlt* (Del Alamo et al, 2022). *AFSample* inferences used various *AF-Multimer* model weights (v2.1.2, v2.2.0, and v2.3.2). In all cases modeling was done with no templates. When using *AF-Multimer v2*.*1*.*2*, modeling was done using 21 max_recycles, with *v2*.*2*.*0* with the default of 3 max_recycles, and with *v2*.*3*.*2* using 9 max_recycles. *AFSample2* inferences used the same variation in *AF-Multimer* model weights as the *AFSample* runs. In all cases inference was run with no templates and 3 max_recycles. Runs with *AFAlt* use *AF-Multimer v2*.*1*.*2*, no templates, 5 max_recycles, and max MSA depths ranging from 32 to 128. Hydrogen atoms were added to files generated by *AFSample, AFSample2*, and *AFAlt* using a custom script which employs the Amber Force Field, analogous to the method employed by the original AF2 manuscript (Jumper et al, 2021). Each of these enhanced sampling methods can be quite aggressive in generating conformational diversity in addition to models that are not physically reasonable: e.g. incorrect amino acid chirality, non-native cis peptide bonds, and other biophysically incorrect features, particularly in the not-well-packed residue segments of the modeled proteins. The most egregious of these physically unreasonable models were identified and removed, as described elsewhere (Spaman et al, manuscript in preparation). The resulting relaxed models were used for further analysis.

CSPs were calculated by comparing chemical shifts between bound and free forms of the complex. CSPs were calculated only for well-defined residues (Snyder & Montelione, 2005; Kirchner & Güntert, 2011) of the receptor protein chain. Well-defined residues were determined using the overlapping region from Dihedral Angle Order Parameter (DAOP) analysis (Hyberts et al, 1992), *CYRANGE* (Kirchner & Güntert, 2011) and *FindCore2* (Snyder et al, 2014), determined by running the PDB NMR conformer ensemble with *Protein Structure Validation Server (PSVS) v2*.*0* (Bhattacharya et al, 2007). In each system, only N- and C-terminal “not well-defined” residues were trimmed, while not-well-defined internal loops were not excluded from analysis.

Chemical shift predictors were used to identify which receptor protein residues have backbone atoms with expected significant ^15^N-^1^H CSPs upon peptide binding. There are multiple state-of-the-art chemical shift predictors including *ShiftX2* (Han et al, 2011), *Sparta+* (Shen & Bax, 2010), and *UCBShift* (Li et al, 2020). Each of these methods has advantages and disadvantages for this application; for instance, when provided with structural homologs with assigned chemical shifts *UCBShift* has been shown to outperform other methods. The analysis presented here uses predicted chemical shifts from *UCBShift* generated using the implementation on NMRBox servers (Maciejewski et al, 2017). However, comparable results were obtained using predicted shifts from other methods. These other results can be accessed via a GitHub repository created for this study (https://github.rpi.edu/RPIBioinformatics/CSP_Rank) or in **Supplementary Figure 1**.

Chemical shifts were acquired from 2D ^15^N-HSQC spectra, providing backbone amide ^1^H and ^15^N CSP data. CSPs were calculated as described elsewhere (Weng et al, 2020):

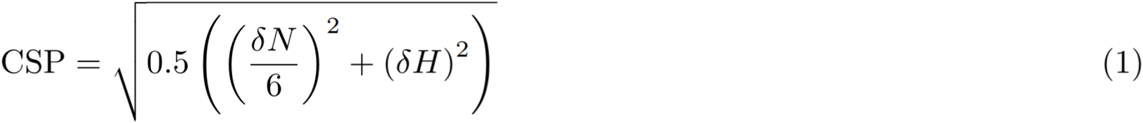

Cutoff values for “significant” CSPs were determined as described previously (Ma et al, 2016; Aiyer et al, 2021). Briefly, significant outlier CSPs were identified using an iterative procedure in which mean and standard deviations of CSPs were computed, all CSPs with values greater than an outlier cutoff (z-score = 3) were removed, mean and standard deviations were then recomputed for the remaining CSPs, and the process was iterated until no CSPs remain that are greater than this outlier cutoff. The subset of CSPs below this final outlier cutoff were used to compute the mean CSP (excluding outliers) and standard deviation. CSPs less than this mean were defined as “insignificant”, those greater than this mean as “significant”. These “significant” and “insignificant” CSPs were then used to generate a ranking metric (described below) to assess the fit of holo structure models to the experimental CSPs.

### Cross validation

Knowledge-based model quality assessments, including *ProCheck* (backbone and all dihedral angles) (Laskowski et al, 1993), Ramachandran plot analysis (Chen et al, 2010), and *MAGE* clashscore analysis (Chen et al, 2010) were done as described elsewhere using the *Protein Structure Validation Server* (PSVS) (Bhattacharya et al, 2007). For systems with available data, the final computational models were cross validated against NMR NOESY and/or RDC data using RPF DP-scores (Huang et al, 2005; Huang et al, 2012) computed using *RPF* and *PDBStat* (Tejero et al, 2013) software implemented in *PSVS*.

## RESULTS

### Dataset of protein-peptide complexes with NMR data

We compiled a dataset including three-dimensional structures of 108 protein-peptide complexes structures and their associated apo and holo chemical shift lists. This dataset contains numerous systems known to involve extensive structural rearrangement upon peptide binding. The dataset includes 22 calmodulin-peptide complexes (Soderling & Stull, 2001), 6 TFIIH-peptide complexes (Rimel & Taatjes, 2018), 9 bromo and extraterminal domain (BET) complexes of BRD3 or BRD4 ET domain bound to polypeptides (Cheung et al, 2021) and 71 other protein-peptide complexes. All the protein-peptide systems in this dataset have 1:1 stoichiometry. The protein receptors range from 34 to 197 residues, and the ligand peptides from 4 to 73 residues. Metrics for DockQ (Basu & Wallner, 2016) and TM (Zhang & Skolnick, 2005) scores, and for backbone atomic RMSDs between the well-defined residues of the protein receptor in the baseline AF2 models relative to the medoid experimental models, are provided in **Supplementary Figure 2**. This dataset, including the PDB and BMRB ID’s, experimental and baseline AF2 atomic coordinates, experimental backbone chemical shifts (apo and holo), and predicted backbone chemical shifts (apo and holo) are available in a publicly accessible GitHub repository (https://github.rpi.edu/RPIBioinformatics/CSP_Rank).

For each system in the protein-peptide complex dataset, five sets of ^15^N-^1^H chemical shift lists were collected from the BMRB archive or generated by chemical shift prediction based on atomic coordinates:

1. apo receptor shifts from BMRB
2. holo receptor shifts from BMRB
3. predicted shifts for baseline AF2 apo structure
4. predicted shifts for baseline AF2 holo structure
5. predicted shifts for PDB holo structure.

Using these data sets, three sets of CSPs were calculated for each complex:

1. CSPs between apo and holo shift chemical shift lists from BMRB
2. CSPs between shifts predicted for baseline AF2 apo and PDB holo structures
3. CSPs between shifts predicted for baseline AF2 apo and baseline AF2 holo structures.

While the focus on this work is on using ^15^N-^1^H data, CSPs were also calculated using backbone ^13^C’ and ^13^Cα-^1^H CSP data (Grzesiek et al, 1996; EvenaÈs et al, 2001, Williamson, 2013). These carbon shift data are less extensive in the BMRB; our database contains only 38 entries with ^15^N-^1^H, ^13^C’, and/or ^13^C-^1^H CSP data. Analysis using these restricted datasets is presented in **Supplementary Figure 3**.

### CSP_Rank_Score

To rank the fit of structure models to CSP data, CSPs were computed for alternative models and compared. For each apo / holo pair, significant and insignificant CSPs are computed as outlined in the Methods section. A confusion matrix (**Table 1**) was generated by comparing predicted vs observed, and significant vs insignificant, CSPs. In computing the confusion matrix, for each observed CSP the absolute value of its z-score is used to increment the corresponding quadrant of the confusion matrix; each increment of TP, FP, FN, and TN is the z-score of the corresponding observed CSP so that CSPs significantly larger (or significantly smaller) than the mean CSP contribute more (or less) to the resulting statistical scores. Hence, there is more penalty applied to a model for missing a very large (significant) CSP than for missing a CSP close to the mean CSP value.

**Table 1.** Confusion matrix used to label CSPs of the protein receptor upon peptide binding as True Positive (TP), False Negative (FN), False Positive (FP), or True Negative (TN). These values are used to calculate statistical measures of the fit of structure models to observed significant CSPs.

|  | Observed Significant CSP | Observed Insignificant CSP |
| --- | --- | --- |
| Predicted Significant CSP | True Positive (TP) | False Negative (FN) |
| Predicted Insignificant CSP | False Positive (FP) | True Negative (TN) |

True Positives (TP) are instances where a CSP is both predicted and observed to be significant (as defined above). False Positives (FP) are instances where a CSP is erroneously predicted to be significant. False Negatives (FN) are instances where a CSP is erroneously predicted as insignificant. True Negatives (TN) are instances where a CSP is both predicted and observed to be insignificant.

Using this confusion matrix, the following information retrieval statistics were computed for each complex and each CSP data set:

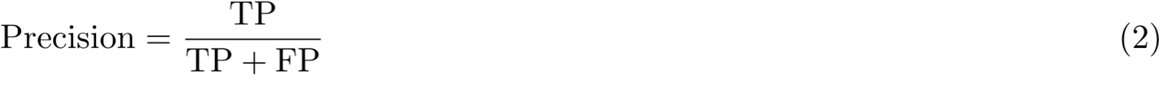

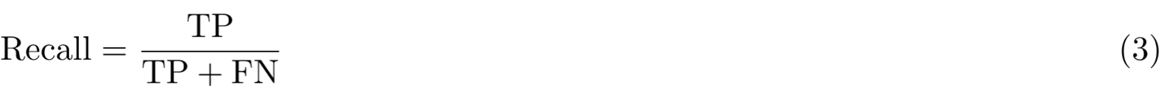

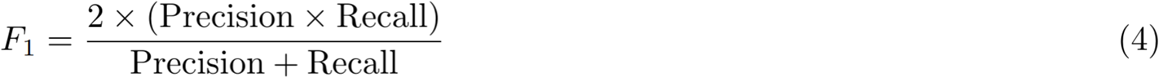

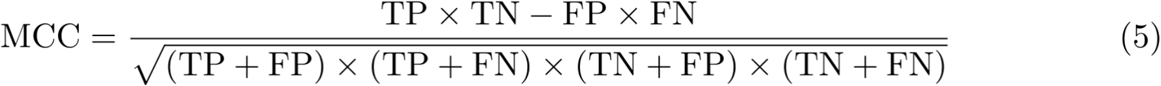

TN’s are not typically factored into an F1 performance score. The Matthew’s Correlation Coefficient (MCC) (Matthews, 1975) is used in information retrieval statistics to account for instances where there is an imbalance confusion matrix; e.g. for systems with commonly encountered in protein-peptide complexes where there are many insignificant CSPs (e.g. for residues distant from the peptide binding site) relative to the number of significant CSPs.

Using these statistics, we define a CSP_Rank_Score as the weighted average of F1 and MCC (Chicco & Jurman, 2020):

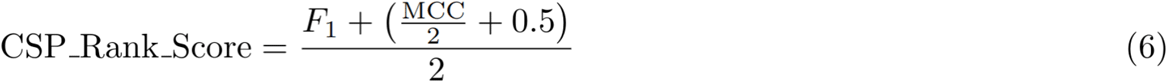

The CSP_Rank_Score is used to compare how well various models of protein-peptide complexes taken from the PDB or predicted by AF2 fit to the CSP data. Comparison plots for F1 and MCC scores across the database for medoid PDB models and top-ranked baseline AF2 models are presented in **Supplementary Figure 4**.

### Performance of AF2 on protein-peptide complexes

Using this collection of 108 protein-peptide complexes for which NMR CSP data are available, the accuracies baseline AF2 to the corresponding medoid PDB model, as assessed by TM Scores, are shown in **Figure 1A**. As was observed in a recent similar study on a database of 96 protein-peptide complexes (Tsaban et al, 2022), baseline AF2 models often have remarkably good agreement with the experimentally determined structures of these complexes. In a few cases, however, AF2 predicts binding sites and poses incongruent with the models deposited in the PDB. The AF2 complex models were also assessed with commonly-used knowledge-based structure quality scores including Procheck G-factors (Laskowski et al, 1993) for both phi/psi backbone dihedral angles or for all backbone and sidechain dihedral angles (**Supplementary Figure 5**), MolProbity clashscores (Chen et al, 2010) (**Supplementary Figure 6**), and Ramachandran plot statistics (Chen et al, 2010) (**Supplementary Figure 7**). These values are reported as Z-scores relative to the corresponding scores in high-resolution X-ray crystal structures (Bhattacharya et al, 2007). Overall, as illustrated in **Supplementary Figures 5-7**, the AF2 models have much better knowledge-based structure quality statistics than the medoid PDB models. A similar observation was made in our studies of AF2 models generated for small proteins and compared with the corresponding experimentally determined structures available in the PDB (Tejero et al, 2022; Li et al, 2023).

**Fig 1.**
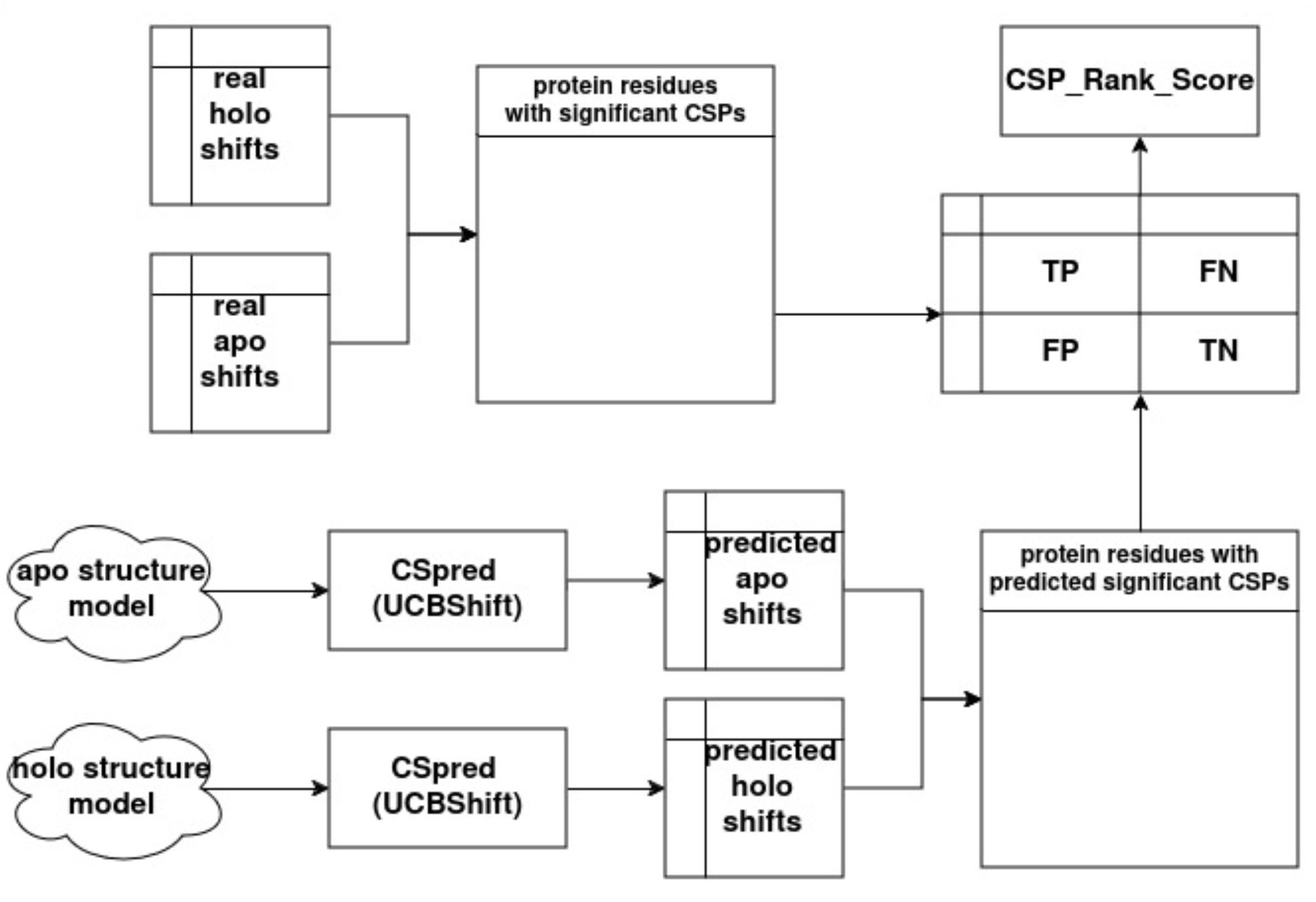
Data flowchart for generating CSP_Rank_Score. Structure models are selected from the PDB archive (medoid model of NMR ensemble) or generated by AF2 and provided to a chemical shift predictor (e.g. UCBshift) to generate a list of residues with predicted significant CSPs (bottom two rows). These predicted significant CSPs are compared to the list of protein residues with experimentally observed (real) significant CSPs shifts (top two rows) to generate a confusion matrix and resulting statistical scores used to compute the CSP_Rank_Score (see **Eqns. 2 - 6**).

Baseline AF2 protein-peptide models also generally perform well when assessed against the experimental CSP data (**Figures 2B, C**). In fact, 67 of the 108 AF2-modeled protein-peptide complexes have better CSP_Rank_Scores than the corresponding medoid PDB model. For each system, chemical shifts were predicted with *UCBShif*t (Li et al, 2020) for the holo medoid model from the deposited NMR ensemble and the apo and holo computational models received from baseline AF2. CSPs which arise from the predicted shifts (bottom half of **Figure 1**) and experimentally observed chemical shifts reported in the BMRB (top half of **Figure 1**) are compared to generate a confusion matrix and CSP_Rank_Score. In each case, the chemical shift lists for apo and holo systems were aligned by applying an offset to the apo data set to minimize the mean difference in chemical shift values between the apo and holo shift lists.

**Fig. 2.**
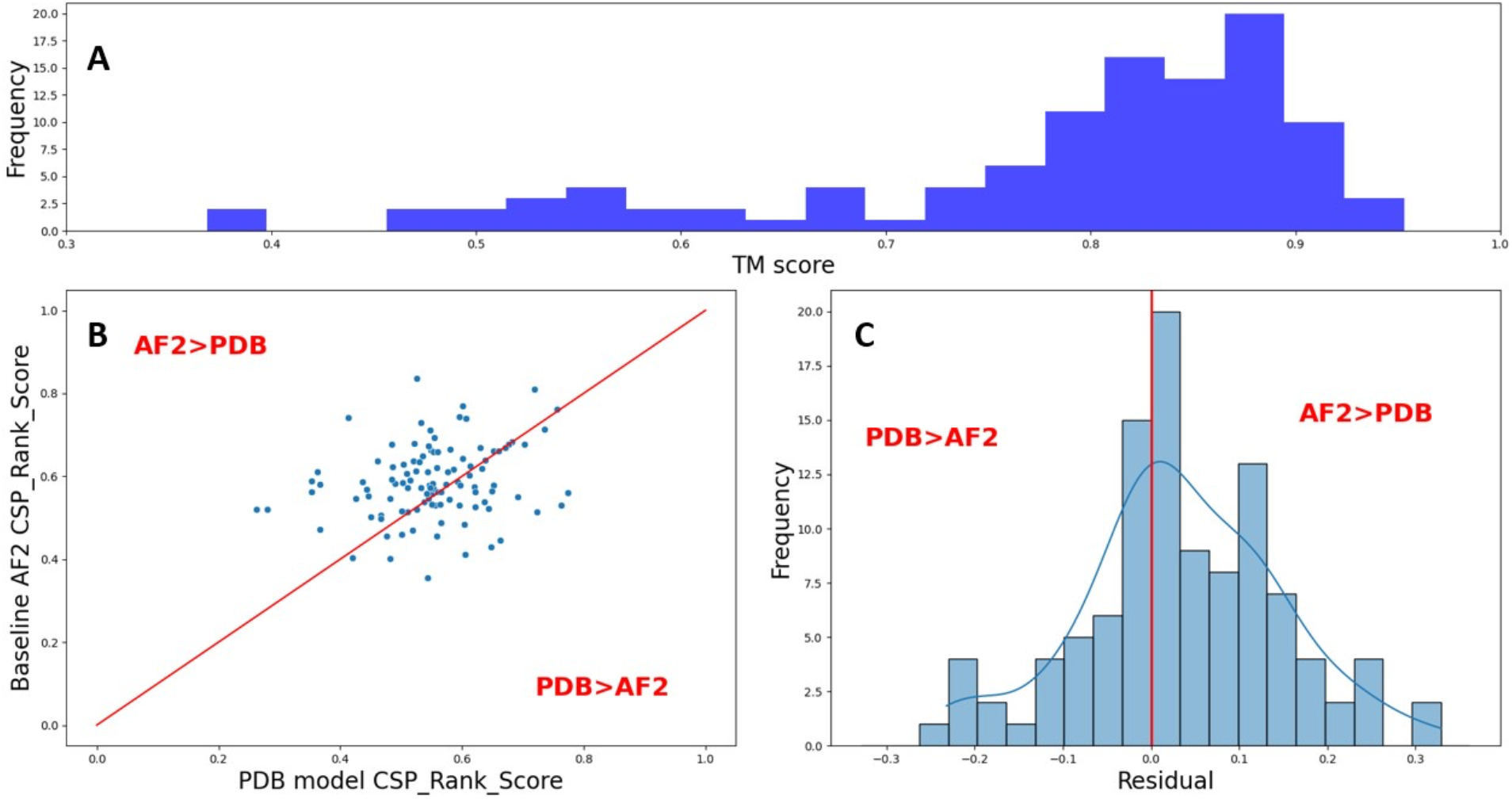
Performance summary of baseline AF2 on protein-peptide dataset. (A) Histogram of TM scores of baseline AF2 models to the medoid model selected from the NMR ensemble deposited in the PDB. (B) Scatter plot of CSP_Rank_Scores for 108 complex models, with y=x line plotted in red. Points above the y=x line denotes systems where the baseline AF2 model fits the CSP data better than the medoid PDB model. (C) Histogram of residuals from the y=x line; when the residual is > 0 then the baseline AF2 model fits the CSP data better than the medoid PDB model. Paired sample t-test p-value = 0.033. During this work, AlphaFold 3 (Abramson et al, 2024) was released. We evaluated performance across our dataset using the AlphaFold 3 server as a structure prediction engine in **Supplementary Figure 8**.

Some representative cases from the study outlined in **Figure 2 are illustrated in Figure 3**. In these plots, we compare the medoid PDB and top-ranked AF2 models, color coding the residues with observed ^15^N-^1^H CSPs on the receptor; with larger CSPs indicated in darker shades of red. A bar plot of CSP vs. residue number is shown at the top of each panel, with coloring of bars indicating the size of the CSP. “Blips” along the x-axis indicate the agreement of the CSPs predicted from the AF2 and/or PDB model with these NMR data; the color coding of these “blips” is explained in **Figure 3 legend**.

The first class of AF2 models, in which the top-ranked baseline AF2 model fits the CSP data better than the medoid PDB model, is illustrated by the case of PDB_IDs 2kpz and 2kfh (**Figure 3A, B**). For 2kpz the CSP_Rank_Score of AF2 model (0.83) is significantly higher than that of the medoid PDB model (0.54); this is also evident from the molecular structure image shown, where in the AF2 model the peptide hugs up closely against residues exhibiting significant chemical shift perturbations and the sites showing the largest CPSs. These interactions are mostly annotated with purple or green blips that indicate good agreement between predicted and observed CSPs in the AF2 model. Model PDB_ID 2kfh (**Figure 3B**) provides a second illustrative example of this case, where the AF2 model (CSP_Rank_Score 0.52) fits the CSP data better than the medoid PDB model (rank score 0.26). Approximately 30% of the AF2 models exhibit better agreement with CSP data than the corresponding PDB model (**Figure 1C** residual > 0.1). As in the previous two examples, in most of these cases there are no yellow blips (indicating the PDB model fits better than the AF2 model) for significant CSPs. A second class of AF2 models is illustrated by the case of PDB_ID 2n7k (**Figure 3C**), where baseline AF2 misplaces the location of the peptide ligand, resulting in a significantly better CSP_Rank_Score for the PDB model (0.61) than for the baseline AF2 model (0.41). Approximately 10% of the AF2 models fall into this class (**Figure 1C** residual < -0.1); here the purely computational AF2 method provides an incorrect model for the protein-peptide complex. The third class of AF2 models, where CSP_Rank_Scores are similar (-0.1 ≤ **Figure 1C** residual ≤ 0.1), and visual inspection does not provide a clear conclusion of which is a better fit to the CSP data, is illustrated by model PDB_ID 7jq8 (**Figure 3D**). Similar performance observed for approximately 60% of the targets. **Figure 3D** also illustrates a case of significant allosteric changes in the receptor upon peptide binding, resulting in CSPs throughout the receptor upon peptide ligand binding and creating significant challenges for any CSP-based docking protocol. *It is remarkable that for ∼ 90% of the protein-peptide complexes in our data set, the CSP_Rank_Scores for AF2 models, generated with no sample-specific experimental data, are similar or better than the CSP_Rank_Scores of the PDB models which are based on CSP and other experimental NMR data*.

**Fig. 3.**
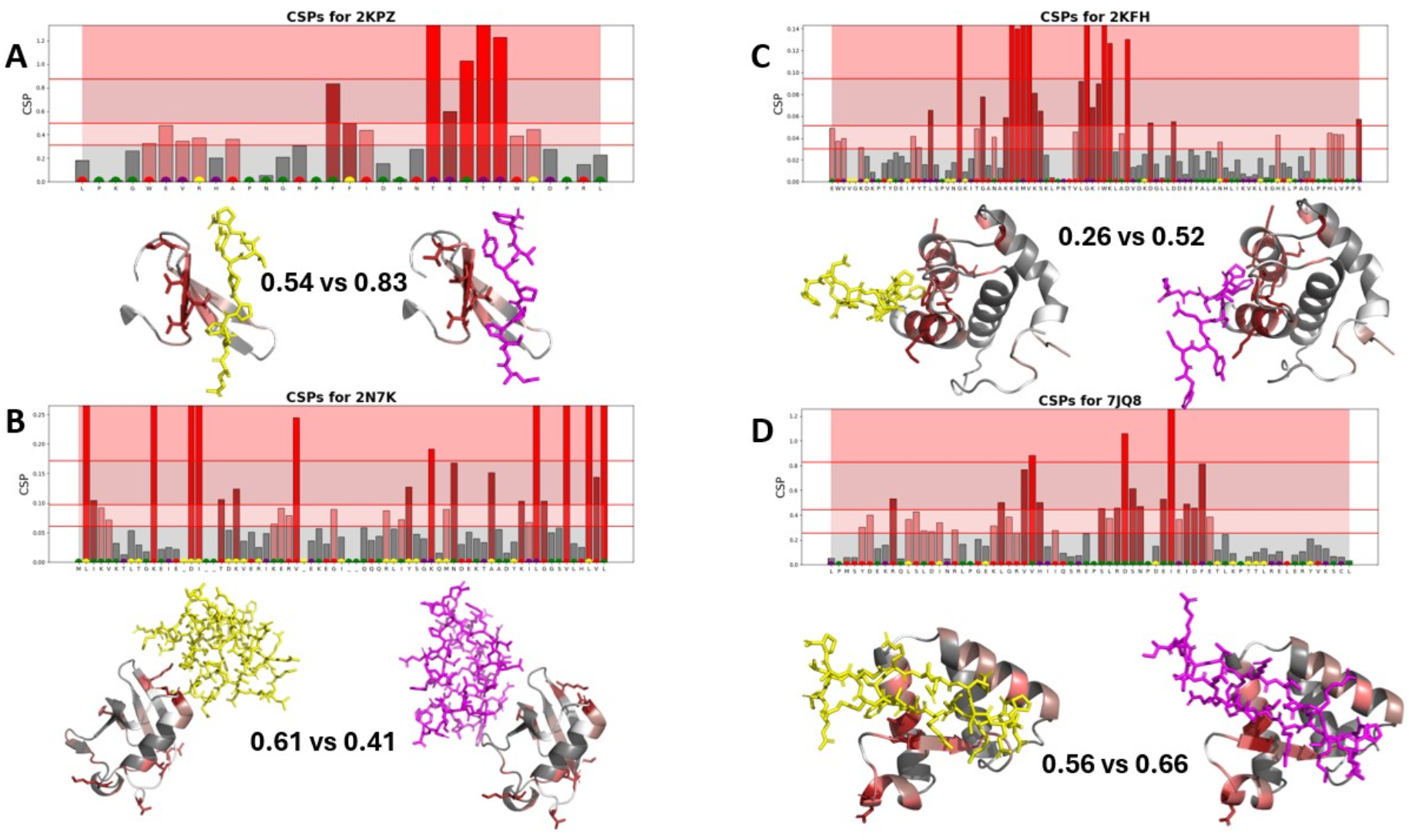
Comparison of predicted and observed CSPs for representative complexes modeled with AF2. In each panel a CSP histogram is provided for each residue in the sequence, with bars colored varying shades of red to reflect the size of the indicated CSP; red bars indicating large “significant” CSPs, and grey bars indicating small “insignificant” CSPs. Colored “blips” at the bottom of the histogram denote the pattern of agreement between the observed CSPs and those predicted using the PDB model or the AF2 model: purple - AF2 model fits the CSP data but the PDB model does not, green - AF2 and PDB models both fit CSP about equally well, yellow - PDB model fits the CSP data but the AF2 model does not, and red - neither AF2 nor PDB model fit the CSP data. Below the histogram is a structural view of the medoid PDB model (left) and rank 1 AF2 model (right) with the backbone cartoon of the protein colored by the significance pattern of CSPs. **(A)** Data for PDB_ID 2kpz, a case where the AF2 model has a much better CSP_Rank_Score. **(B)** Data for PDB_ID = 2kfh, another case where AF2 achieves a better CSP_Rank_Score by docking closer to residues with significant CSPs. **(C)** Data for PDB_ID=2n7k, a case where AF2 misplaces the docking of the peptide ligand, which results in a better CSP_Rank_Scorefor the PDB model. **(D)** Data for PDB_ID 7jq8, a case with similar CSP_Rank_Scores for the two models, and many allosteric CSPs throughout the receptor due to peptide binding. In all cases, the top-ranked baseline AF2 model and medoid conformer from the PDB NMR ensemble are used to predict CSPs, which are then compared to the experimental CSP data to calculate a CSP_Rank_Score.

### Enhanced sampling with AF2

We reasoned that enhanced sampling (ES) with AF2 could generate a diverse collection of models, among which would be models that fit even better to the CSP data than models generated with baseline AF2. For several systems, we explored this hypothesis using three different ES methods, *AFSample* (Johansson-Åkhe & Wallner, 2022), *AFSample2* (Kalakoti & Wallner, 2024), and *AFAlt* (Del Alamo et al, 2022). For this work we used published protocols for the ES methods, which determined the number of models generated with each method. The distributions of structures generated by ES methods were clustered, each of the resulting models was annotated using a Bayesian CSP_Rank_Score (outlined below), and the top-scoring models from each cluster were used to generate an ensemble of complex models representing the uncertainty of the resulting AF2-NMR model.

### Bayesian model selection score and model selection

For each model generated by the enhanced AF2 sampling protocol, we estimated a likelihood of the model from an unnormalized Bayes equation:

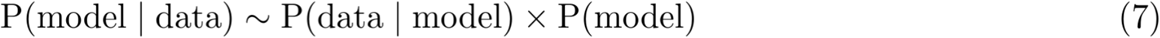

where

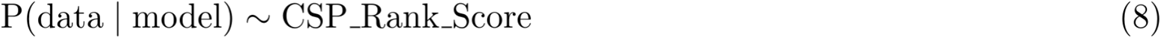

describes how well the data is explained by the model, and the Bayesian prior is

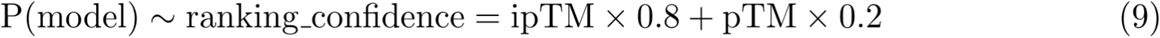

Here, ipTM and pTM are AF2 reliability scores (Jumper et al, 2021) for the corresponding model; this prior is the ranking confidence score defined by Wallner (Wallner, 2023b).

To rank models from the ES protocols, we use the following equation

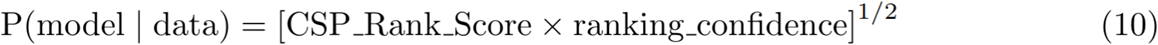

### AF2 - NMR models of protein complexes

To characterize the conformational heterogeneity sampled by *AFSample, AFSample2, and AFAlt*, residues of the receptor proteins that are well-defined (described in Methods) across the resulting ensemble were superposed using the medoid model from the ensemble deposited in the PDB as a reference. The Bio.3d R package (Grant et al, 2006) was then used to perform a principal component analysis (PCA) by aligning the coordinates of the receptor (excluding the not-well defined N- and C-terminal tails) and using the variation in coordinates of the peptide ligand for PCA. The first three principal components of the resulting PCA are used to identify and remove outlier structures, using an algorithm that calculates the Mahalanobis distance (Mahalanobis, 1936; Venables & Ripley, 2002; Brereton & Lloyd, 2016) for each point in the PCA space and excludes points beyond a defined statistical threshold (97.5th percentile of Chi-squared distribution with 3 degrees of freedom). After iteratively removing outliers detected in PCA space, the coordinates are analyzed by t-SNE (van der Maaten & Hinton, 2008, van der Maaten L, 2014; Krijthe, 2015) and UMAP (McInnes et al, 2018) non-linear dimensional reduction. K-means hierarchical clustering was then performed in both t-SNE and UMAP space. The single top-scoring model, in terms of their P(model|data), from each cluster was selected and incorporated into an ensemble of structures representing the model of the complex and its uncertainty. We refer to the resulting ensemble as the *AF-NMR model*, in which ES with AF is combined with experimental data to generate an ensemble representation of the set of AF models which best fit the experimental CSP data.

A detailed illustrative example of the *CSP_Rank* protocol for protein-peptide complexes is presented in **Figure 4** for the complex formed between the extra-terminal domain (ET) of the bromodomain and ET domain protein (BET protein) (BRD3) and the C-terminal polypeptide segment of the murine leukemia virus integrase protein (TP), PDB_ID 7jq8, a structure from our own lab for which we have access to the raw experimental data (Aiyer et al, 2021). This system involves a disorder-to-order transition upon binding of the 23-residue TP to the receptor ET protein. ET-TP is a particularly challenging system because the ET protein receptor itself also becomes more ordered upon binding, as a disordered loop forms a three-stranded beta-sheet with a hairpin conformation of the protein, and there are allosteric changes of ET upon complex formation resulting in small CSPs throughout the ET receptor structure. In this case, baseline AF2 has a better fit to the CSP data than the PDB medoid structure based on extensive experimental NOESY data (CSP_Rank 0.65 vs 0.56) (**Figure 3D**). **Figure 4A-D** illustrates the diversity of models generated with various ES protocols with AF2. Selection of the models from each cluster with highest CSP Bayesian model selection score results in the ensemble shown in **Figure 4E**. In this model, the interactions between the TP and ET are extensive and well-defined. The model has excellent knowledge-based structure quality scores; *viz* Procheck phi/psi Z-score +1.14, ProCheck all dihedral Z score +2.07; MolProbity clashscore Z-score +0.15, with 100% of dihedral angles in the most favored regions of the Ramachandran map (**Supplementary Table 1**), that are better than those of either the PDB (**Supplementary Table 3**) or baseline AF2 models, and structural protein-peptide interaction features that are very consistent with those determined by the experimental NMR structure (Aiyer et al, 2021) (**Table 2; Supplementary Figure 9**). Interestingly, the conformational variations (backbone atomic coordinate RMSFs) across this AF-NMR ensemble are highly correlated with the residue-specific pLDDT score averaged across the individual AF2 models of the ensemble (**Figure 4F**), which is consistent with our similar observations on other systems (Huang & Montelione, 2024; Spaman et al, manuscript in preparation).

**Fig. 4:**
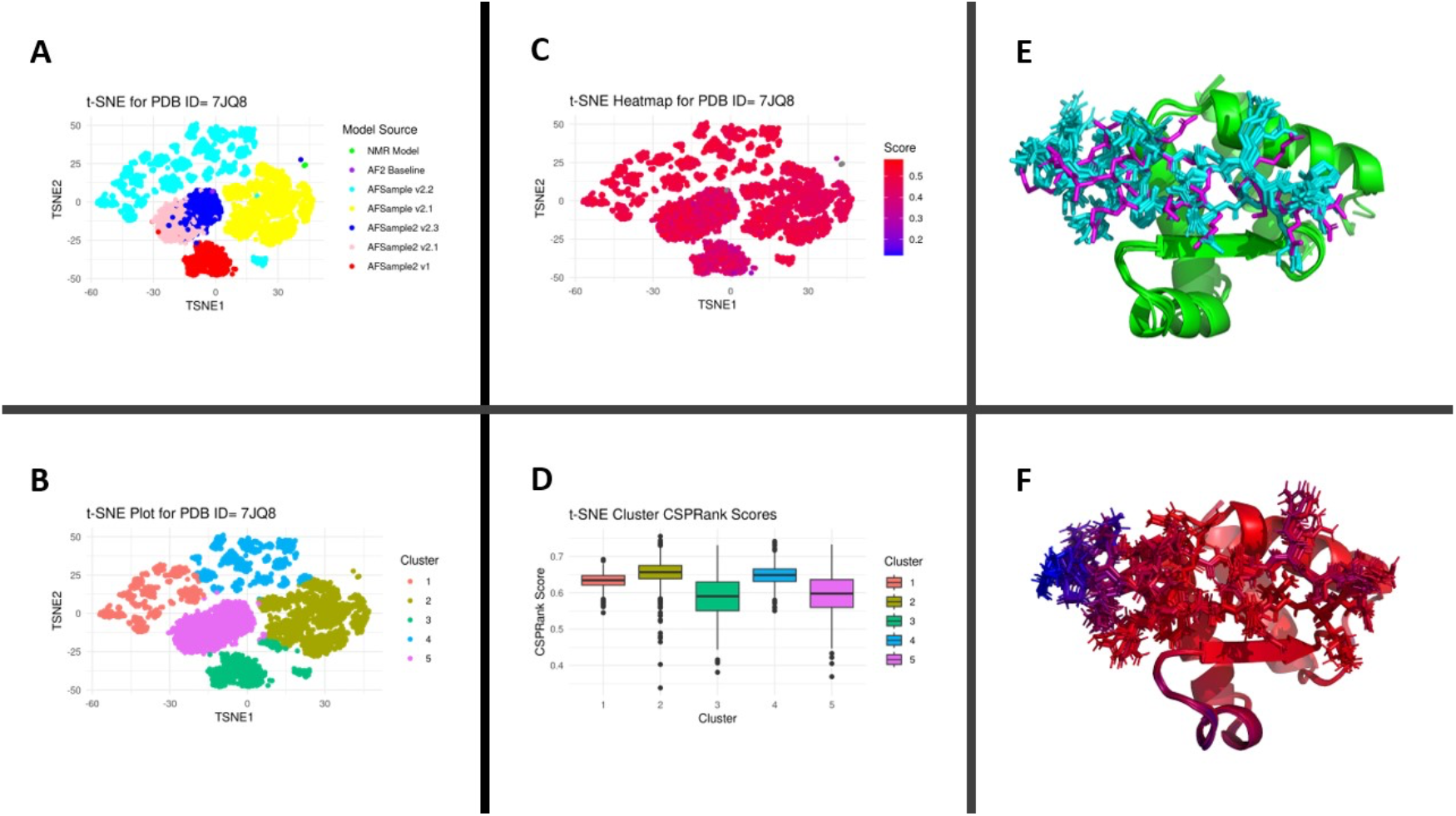
Enhanced sampling for IN TP:ET complex. (**A)**. Plots of first two TSNE PCA dimensions demonstrating conformational space explored by ES methods AFSample and AFAlt. Different ES sampling protocols provide different models of the complex. (**B**) Clusters extracted from K-means hierarchical clustering. **(C)** Heatmap of P(model|data) overlaid on TSNE analysis. (D) Boxplots of CSP_Rank_Scores from each TSNE cluster. **(E)** Ensemble of the models which have the highest CSP_Rank_Score from each TSNE or UMAP cluster, colored by chain, where the receptor protein is green, and the ligand peptide is blue. **(F)** Ensemble depicted in 4E; the color encodes the residue-specific pLDDT score averaged across the AF2 models of the ensemble (red - high; blue - low).

**Table 2.** Cross-Validation RPF results for 7JQ8 and 7JYN. Table summarizing RPF statistics and CSP_Rank_score for each type of ensemble. AF-NMR ensembles were derived using **Eqn 8** as the selection criteria (CSP-based selection), AF-NMR* ensembles were derived using **Eqn. 11** as the selection criteria (CSP / NOE –based selection). RPF statistics are reported as the average of values generated from individual models in the ensemble. CSP_Rank_Scores are reported according to the value of the medoid model in each ensemble. The modified selection criteria for AF-NMR* ensembles result in better agreement with NOE data than a selection metric unbiased by NOE observables; in both cases this new selection metric does not substantially impact the value of the medoid model CSP_Rank_Score and only changes the atomic coordinates slightly. **Supplementary Figure 9** provides a Double Recall plot (Huang, Montelione 2024) for 7JQ8 highlighting the unique peaks satisfied by the AF-NMR* ensemble for a specific Trp residue. **Supplementary Figure 20** provides a Double Recall plot for 7JYN. These Double Recall plots demonstrate how the AF-NMR* models are consistent with some key interfacial NOESY peaks not explained by the PDB models. **Supplementary Figure 9** contains t-SNE clustering of 7JYN AF-NMR ensemble like those shown in **Figure 4** for 7JQ8. **Supplementary Tables S9-14** provide PSVS structure quality reports for each of the complexes included in Table 2.

|  | PDB |  | AF-NMR |  | AF-NMR* |  |
| --- | --- | --- | --- | --- | --- | --- |
|  | 7JQ8 | 7JYN | 7JQ8 | 7JYN | 7JQ8 | 7JYN |
| RPF Recall | 0.88 | 0.92 | 0.88 | 0.90 | 0.88 | 0.90 |
| RPF Precision | 0.84 | 0.75 | 0.85 | 0.75 | 0.86 | 0.76 |
| RPF DP-Score | 0.646 | 0.698 | 0.675 | 0.659 | 0.685 | 0.680 |
| CSP_Rank_Score | 0.56 | 0.49 | 0.74 | 0.77 | 0.73 | 0.78 |

The ES AF2-NMR protocol outlined in **Figure 4** was applied to 17 complexes from the protein-peptide database. In all cases, the medoid model selected from the ES ensemble (hereafter denoted as AF-NMR ensemble) exhibits excellent knowledge-based structure quality scores and better fit to the CSP data than the baseline AF2 model (**Supplementary Tables S1-S14**). For 14 of these systems, the medoid AF-NMR model is also a better fit to the CSP data than the medoid model of each ensemble deposited in the PDB, which are based on extensive additional NMR data (**Supplementary Figure 10**). Some examples comparing the medoid model of the AF-NMR ensemble with the top-ranked baseline AF2 model are shown in **Figure 5**. Also included here is the ET-TP complex (**Figure 5D**) used as an illustrative example of the AF2-NMR protocol in **Figure 4D**.

**Fig. 5.**
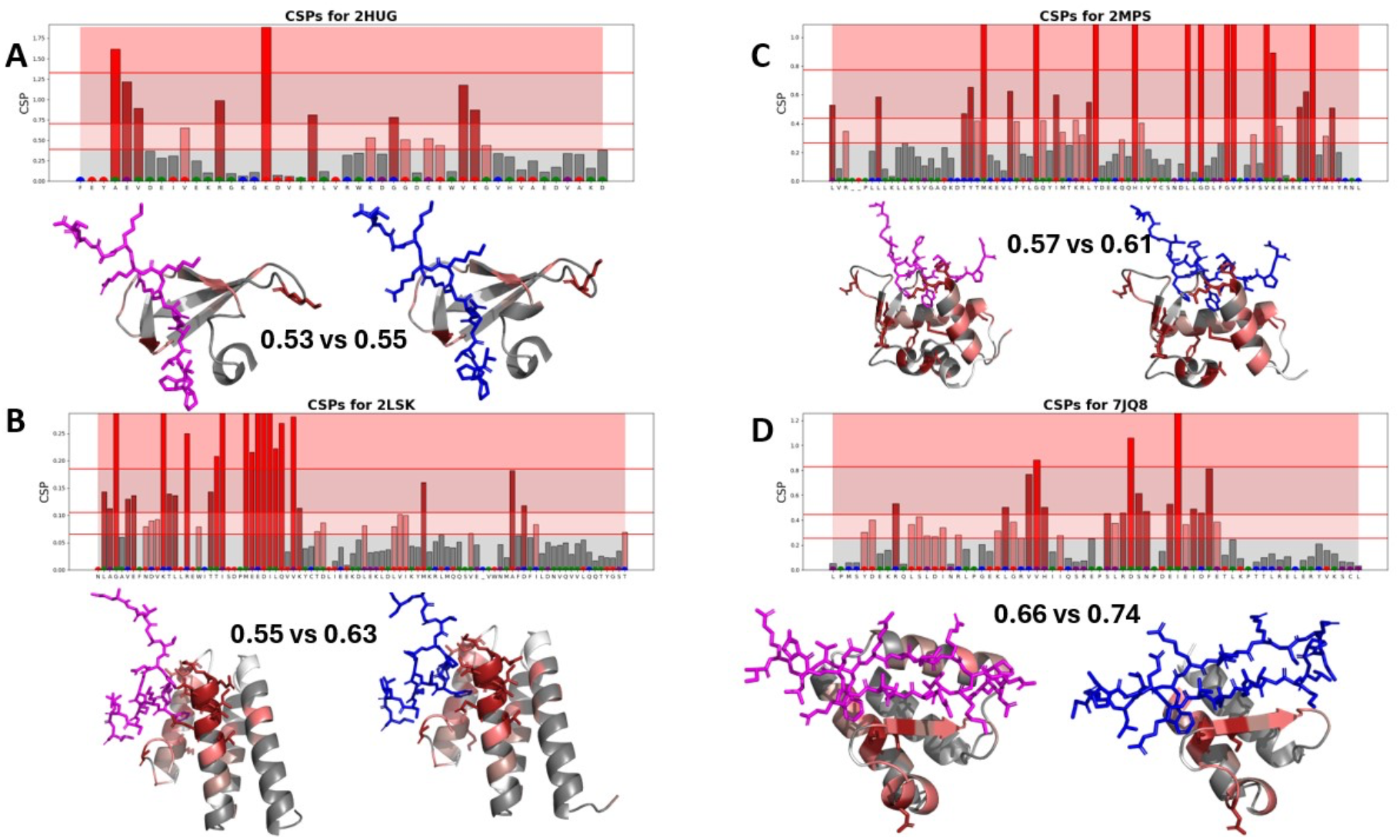
Improvement in CSP_Rank_Scores with Enhanced Sampling. Following the convention from **Figure 3**, in each panel a CSP bar plot is provided for each residue in the sequence, with bars colored varying shades of red to reflect the size of the indicated CSP. Colored blips at the bottom of the histogram denote the pattern of agreement between the observed CSPs and those predicted using the medoid PDB model or the medoid model from the enhanced sampling ensemble as defined in **Figure 4** legend. Below the bar plot is a structural view of the top-ranked baseline AF2 model (left) and the medoid AF-NMR model (right) with the backbone cartoon of the protein colored by the significance pattern of CSPs. In each of these cases, the ES protocol results in small adjustments to the orientation of the binding peptide ligand which enables a better fit to the NMR CSP data. Across 17 systems tested, the average improvement in CSP_Rank_Score between baseline AF2 model and the medoid AF-NMR model is 0.08. For each of these 17 systems, an ensemble was generated for which the medoid model has a better CSP_Rank_Score than the baseline AF2 model (**Supplementary Figures. 11-19**).

### Cross validation against other experimental data

Although most of the protein – peptide PDB structures used in this study were generated using distance restraints derived from NMR data, we did not attempt to validate the AF NMR models against such distance restraints because the restraint themselves are derived information with inherent (sometimes significant) inaccuracies that result from ambiguities in defining them from the experimental NMR data. Instead, for cross validation we used only systems for which *NOESY peak list data* is available in the PDB or BioMagResDB. Unfortunately, even though such data is invaluable for validation of models and methods, most groups do not provide such “raw” experimental NMR data.

To perform this cross-validation, we rely upon previous work (Huang et al, 2005) which has established a statistical platform, *RPF*, for assessing the fit of models to NOESY peak lists. The *RPF* statistics provide data for Recall (percentage of NOESY peaks that can be explained by the input query structure(s) with a distance cut-off ≤ 5 Å), Precision (percentage of ^1^H–^1^H distances ≤ 5 Å calculated from the query structure that are observed in the NOESY data), F-score, which is the mean of Recall and Precision, and a Discrimination Power (DP) score, which scales the F-score based of the F-score that would be obtained for a random-coil model. The DP score is a value between 0 and 1 assessing the agreement of the model and NOESY peak list data. For two protein-peptide complexes for which NOESY peak list data are available, 7JQ8 (BET ET domain bound to the “tail-peptide” corresponding to the C-terminal segment of murine Moloney leukemia virus integrase) and 7JYN (BET ET domain bound to a peptide fragment of the chromatin-associated host protein NSD3), we assessed the fit of AF-NMR ensembles to the NOESY peak lists using RPF-DP statistics (**Table 2**). For 7JQ8, the average of the individual model DP-scores is slightly better for the for the AF-NMR ensemble than the experimental NMR ensemble deposited in the PDB (0.675 vs 0.646). For 7JYN, the average of the individual model DP-score for the AF-NMR ensemble is somewhat lower than the ensemble deposited in the PDB (0.659 vs 0.698). *However, in both cases the DP scores for the AF2-NMR models determined using only CSP are similar to those of the experimental PDB models determined using these same NOESY data as driving restraints*. Also in both cases, the AF-NMR structures have significantly better agreement with the CSP data, and higher CSP_Rank_Scores than the conventional restraint-based PDB structures (**Table 2**).

### Bayesian-based joint selection protocol

We also explored a *joint selection protocol*, analogous to methods proposed elsewhere to select models based on both NOESY and chemical shift data (Huang & Montelione, 2024). To create a joint selection score, **Eqn. 8** is redefined as:

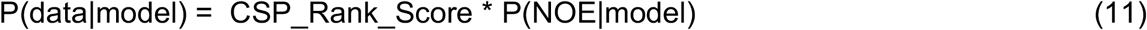

where

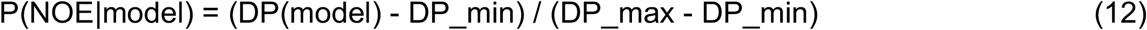

and DP_min and DP_max are the lowest and highest, respectively, DP scores observed across all the ES AF2 models, and DP scores are computed from the back-calculated and observed NOESY peak list using the RPF program (Huang et al, 2005, Huang et al, 2012) available from the ASDP software package (Huang et al, 2015). We refer to these models, jointly selected using both CSP and NOE data, as AF-NMR*.

Selecting models using this joint selection score resulted in models with slightly improved fit to the NOE data without a significant change in the fit to CSP data (7JQ8 AF-NMR* DP-score = 0.685, CSP_Rank_Score = 0.73; 7JYN AF-NMR* DP-score = 0.680, CSP_Rank_Sore = 0.78) (**Table 2**). *These results demonstrate a generalization of our Bayesian selection score to include both CSP and NOESY data, and the potential to expand the score to include additional interfacial experimental data in the AF-NMR protocol where available*.

## DISCUSSION

This study aimed to ascertain how emerging ML-based structure prediction methods can be used to validate or improve upon models of protein-peptide complexes which have experimentally acquired NMR CSP data, and to develop protocols for combining AF2 modeling with CSP for determing 3D structures of protein-peptide complexes. We have developed a scoring function based on NMR CSPs and demonstrated its effectiveness in ranking models from computational methods like AlphaFold2 and AFSample. Our results underscore the value of integrating accessible experimental data, such as NMR CSPs, into computational modeling to enhance their reliability. Approaches that combine computational and experimental techniques have the potential for more accurate and reliable protein structure predictions and may also facilitate modeling conformational dynamics and the relative populations of different conformational states (Huang & Montelione, 2024).

In this study, we explored the concept of model selection from a collection of models generated by enhanced sampling with AF2, using a Bayesian score to rank the fit of protein-peptide structures to experimentally observed NMR chemical shift perturbations. This approach addresses a significant need in the field for validating computational predictions with realistic experimental data. AFSample (Wallner, 2023a), AFSample2 (Kalakoti & Wallner, 2024), AFAlt (Del Alamo et al, 2022), and similar methods (Bryant & Noé, 2024; Wayment-Steele et al, 2024; Stein & McHaourab, 2022) are the current state-of-the-art in predicting diverse structural conformations by employing combinations of MSA subsampling, MSA column masking, neural network dropout, and multiple model training weights. Other methods including molecular dynamics simulations (Mondal, et al 2023) and diffusion-based sampling (Abramson et al, 2024) also may be useful to generate physically-reasonable models which can be ranked using experimental data.

Rather than using the NMR data to drive the AF2 modeling, we allow the ES AF2 modeling to generate unrestrained models and then select those models that best fit the data. This same approach, utilizing a Bayesian framework to combine NOESY and dynamic structural data based on chemical shifts, was also explored in our recent study modeling multiple conformational states the enzyme Gaussia luciferase (Huang & Montelione, 2024) with ES AF2. In both of these approaches, we opted to integrate AF2 modeling with NMR data by conformational selection, rather than using a conventional restraint-based implementation with experimental restraints as input into the AF2 modeling, in order to circumvent the many ambiguities and inaccuracies associated with interpreting NMR data as conformational distance restraints.

As representative examples of cross validation of model predictions using the *CSP_Rank* protocol, we illustrated two cases where generating diverse conformational ensembles with AFSample and AFSample2 and ranking them against NMR CSPs and NMR NOEs resulted in better fits to CSP data and similar fits to NOESY (DP score) data than either the baseline AlphaFold2-Multimer or the experimental PDB structure models. Thus far, 17 systems have been tested with the *CSP_Rank* protocol. Most of these systems result in structural ensembles with better agreement with the CSP data than either the baseline AF2 model or the model deposited in the PDB (**Supplementary Figure 10**).

The primary AF-NMR method described here uses ^15^N-^1^H CSP data for model selection from the ES AF2 ensembles. We also demonstrated the joint use of CSP and NOESY data. Conformer selection from ES AF2 ensembles with NOESY and chemical shift data has also been demonstrated in studies of the conformational dynamics of the enzyme Gaussia luciferase (Huang & Montelione, 2024). However, it is also possible to use other NMR data for such conformer selection, including chemical shift data indicating conformational flexibility (Huang & Montelione, 2024), or data from residual dipolar coupling (RDC) (Chiliveri et al, 2021), nuclear relaxation dispersion, and/or paramagnetic relaxation experiments, other kinds of chemical shift data including chemical shifts determined by chemical exchange by saturation transfer (CEST) data, and other types of NMR, chemical crosslink, or biophysical data. Validating structures generated by ES methods with experimental data provides an important novel approach for accurate analysis of biophysically relevant patterns of structural heterogeneity (Huang & Montelione, 2024).

Although we and others (Ko & Lee, 2021; Tsaban et al, 2022) have observed that AF2 alone does a remarkably good job in predicting the structures of protein-peptide complexes, a significant fraction of these models are incorrect, and there is no way to be certain about the resulting model accuracy without some experimental data. In a recent study of the accuracy of AF2 in modeling of the structures of 96 protein-peptide complexes, 37% had accuracies of < 2.5 Å; 63% had poorer accuracy (Tsaban et al, 2022). Hence, there remains a need for experimental validation and refinement of these assembly models. This integrative AF2-NMR approach provides accurate models of protein-peptide complexes using only ^15^N-^1^H CSP data, and is easily extended to include ^13^C CSP, NOESY, RDC, or other kinds of interfacial experimental data. As only backbone resonance assignments for the receptor are required, there is also no need to produce isotope-enriched polypeptides, which can be both expensive and challenging. Overall, the AF2-NMR approach has the potential to significantly advance the field of protein-peptide complex modeling, providing more accurate models that can better inform biological research and therapeutic development.

## Author Contributions

TLB and GTM jointly conceived this study and analyzed and interpreted data. TLB wrote computer codes, generated graphics, and organized the GitHub Data Repository for this paper. Both authors contributed to writing and editing the manuscript.

## Declaration of Interests

GTM is a founder of Nexomics Biosciences, Inc. This does not represent a conflict of interest in this study.

## Acknowledgements

We thank Dr. Davide Sala for providing scripts for running *AFAlt*, T. Acton, A. De Falco, K. Fraga, A. Gaur, R. Greene-Cramer, Y.J. Huang, T.A. Ramelot, B. Shurina, L. Spaman, G.V.T. Swapna, and R. Tejero for helpful discussions and comments on the manuscript, and S. Collen for computer system administration support. We also acknowledge access to the RPI Center for Computational Innovations (CCI) computing infrastructure. This work was financially supported by National Institutes of Health NIGMS grant R35 GM141818 (to G.T.M.) and by the Rensselaer Polytechnic Institute (RPI) Bio-computing and Bio-informatics Constellation Chair Fund. TLB is supported by a NIGMS Biomolecular Science Engineering Training Program T32GM141865.

## Code Availability

Software together with input and output datasets for the examples demonstrated in this paper are available https://github.rpi.edu/RPIBioinformatics/CSP_Rank.

## Supplemental Information

Supplementary material includes 14 Supplementary Tables and 20 Supplementary Figures.

## Abbreviations

AF2: *AlphaFold2 Multimer*
BMRB: Biological Magnetic Resonance Data Bank
CEST: Chemical Exchange by Saturation Transfer
cryoEM: cryogenic Electron Microscopy
CSP: Chemical Shift Perturbation NMR data
ES: Enhanced Sampling
ESM: *Evolutionary Scale Modeling*
LDDT: Local-Distance Difference Test
ML: Machine Learning
MSA: multiple sequence alignment
NMR: Nuclear Magnetic Resonance
NOE: Nuclear Overhauser Effect
NOESY: NOE SpectroscopY
PDB: Protein Data Bank
PSVS: *Protein Structure Validation Server*
pLDDT: predicted LDDT, a model confidence score predicted from ML
RDC: Residual Dipolar Coupling.

## Notes

https://github.rpi.edu/RPIBioinformatics/CSP_Rank

