## Supplementary Material for "Integrative Modeling of Protein-Polypeptide Complexes by Bayesian Model Selection using AlphaFold and NMR Chemical Shift Perturbation Data"

### Table of Contents

**Supplementary Figures**
**Fig. S1** Performance across the dataset using alternative chemical shift predictors.
**Fig. S2** Structure similarity scores between baseline AF2 models and medoid PDB models.
**Fig. S3** Performance across the dataset using alternative calculations for CSPs.
**Supplementary Note 1**. Alternative calculations for CSP using Carbon chemical shifts
**Fig. S4** Performance across the dataset for F1, MCC and CSP_Rank_Score statistics.
**Fig. S5** Knowledge based statistical scores between baseline AF2 and medoid NMR models.
**Fig. S6** Knowledge based Molprobity clashscores between baseline AF2 and medoid NMR models.
**Fig. S7** Knowledge based Ramachandran dihedral angle violations between baseline AF2 and medoid NMR models.
**Fig. S8** Performance across the dataset using AF3.
**Fig. S9** RPF Double Recall analysis for 7JQ8.
**Fig. S10** Improvements to baseline AF2 CSP Rank Score using the *CSP_Rank* protocol.
**Fig. S11** t-SNE clustering analysis and AF-NMR ensemble for 7JYN.
**Fig. S12** t-SNE clustering analysis and AF-NMR ensemble for 2MNU.
**Fig. S13** t-SNE clustering analysis and AF-NMR ensemble for 2JW1.
**Fig. S14** t-SNE clustering analysis and AF-NMR ensemble for 2MPS.
**Fig. S15** t-SNE clustering analysis and AF-NMR ensemble for 2LSK.
**Fig. S16** t-SNE clustering analysis and AF-NMR ensemble for 6H8C.
**Fig. S17** t-SNE clustering analysis and AF-NMR ensemble for 5TP6.

**Fig. S18** t-SNE clustering analysis and AF-NMR ensemble for 2KNE.
**Fig. S19** t-SNE clustering analysis and AF-NMR ensemble for 6BNH.
**Fig. S20** RPF Double Recall analysis for 7JYN.

**Supplementary Tables**

**Table S1** 2MNU AF-NMR Ensemble PSVS structure quality factors.
**Table S2** 2JW1 AF-NMR Ensemble PSVS structure quality factors.
**Table S3** 2MPS AF-NMR Ensemble PSVS structure quality factors.
**Table S4** 2LSK AF-NMR Ensemble PSVS structure quality factors.
**Table S5** 6H8C AF-NMR Ensemble PSVS structure quality factors.
**Table S6** 5TP6 AF-NMR Ensemble PSVS structure quality factors.
**Table S7** 2KNE AF-NMR Ensemble PSVS structure quality factors.
**Table S8** 6BNH AF-NMR Ensemble PSVS structure quality factors.
**Table S9** 7JQ8 AF-NMR Ensemble PSVS structure quality factors.
**Table S10** 7JQ8 AF-NMR* (joint fitting to CSP and NOESY data) Ensemble PSVS structure quality factors.
**Table S11** 7JQ8 NMR Ensemble (directly from PDB) PSVS structure quality factors.
**Table S12** 7JYN AF-NMR Ensemble PSVS structure quality factors.
**Table S13** 7JYN AF-NMR* (joint fitting to CSP and NOESY data) Ensemble PSVS structure quality factors.
**Table S14** 7JYN NMR (directly from PDB) Ensemble PSVS structure quality factors.

**SI References**

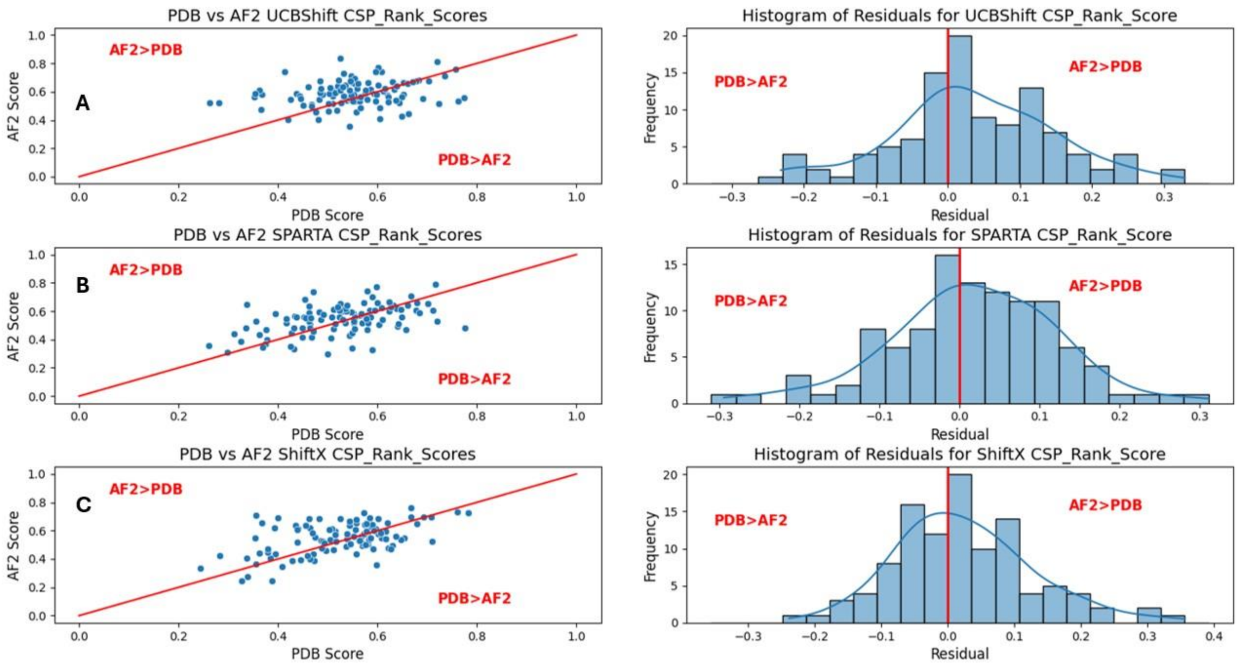


**Fig. S1.** Performance across the dataset using alternative chemical shift predictors. A) Chemical shifts predicted using SPARTA+ (Shen & Bax, 2010). B) Chemical shifts predicted using ShiftX2 (Han et al, 2011) C) Chemical shifts are predicted using SPARTA+, ShiftX2, and UCBShift (Li et al, 2020) and the mean value for each shift is used as a ‘consensus’ shift to generate predicted CSPs. These results demonstrate that regardless of the chemical shift predictor used, our general findings remain the same: baseline AF2 protein-peptide models tend to achieve similar or superior fit to CSP data compared to the medoid model deposited in the PDB.

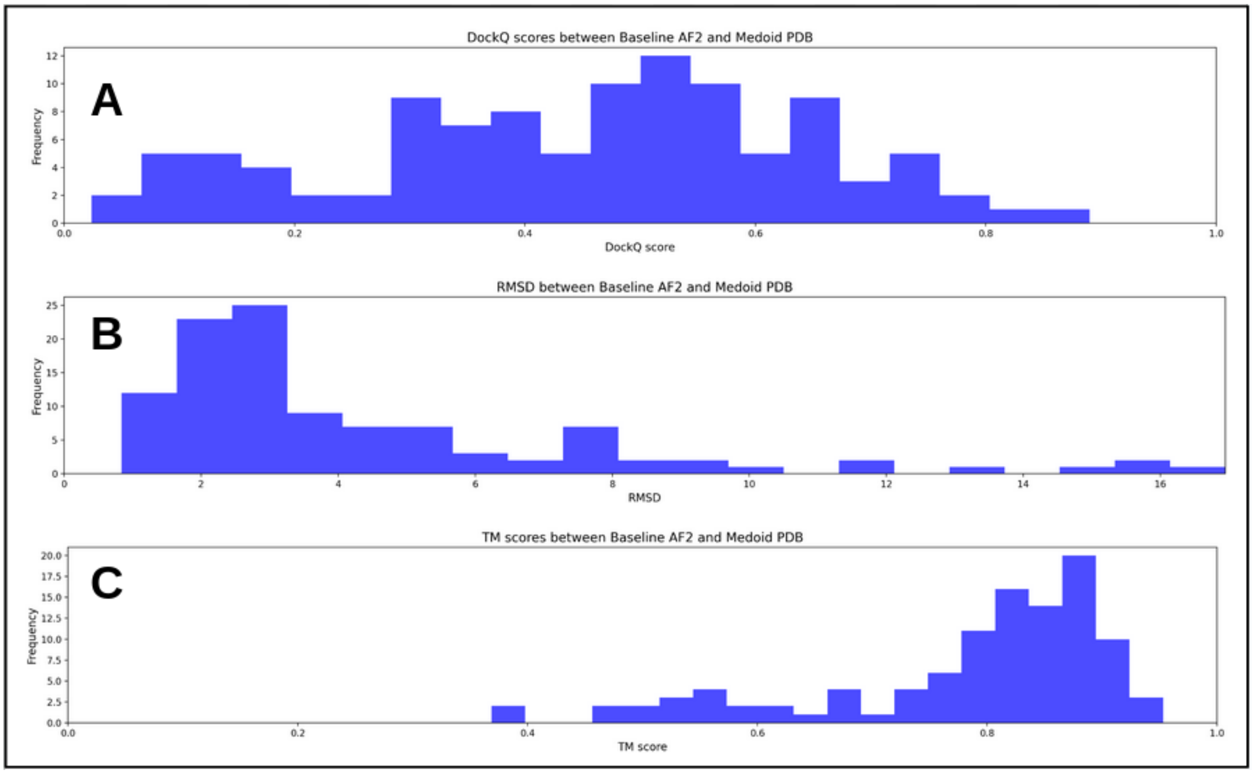


**Fig. S2.** Structure similarity scores between baseline AF2 models and medoid PDB models. A) DockQ (Basu & Wallner, 2016) comparisons. B) RMSD comparisons. C) TM scores retrieved from TM-align (Zhang & Skolnick, 2005). All comparisons involve a structure alignment. The components of each structure used for alignment are the 'well-defined residues' of the protein receptor. Well-defined residues were determined using the overlapping region from DAOP (Kirchner & Güntert, 2011) and FindCore2 (Snyder et al, 2014), determined by running the PDB ensemble with PSVS v2.0 (Bhattacharya et al, 2007). In each system, only N- and C-terminal "not well-defined" residues were trimmed. All residues were used to calculate the statistic after structural alignment, regardless of the status of the residue belonging to the set of well-defined residues.

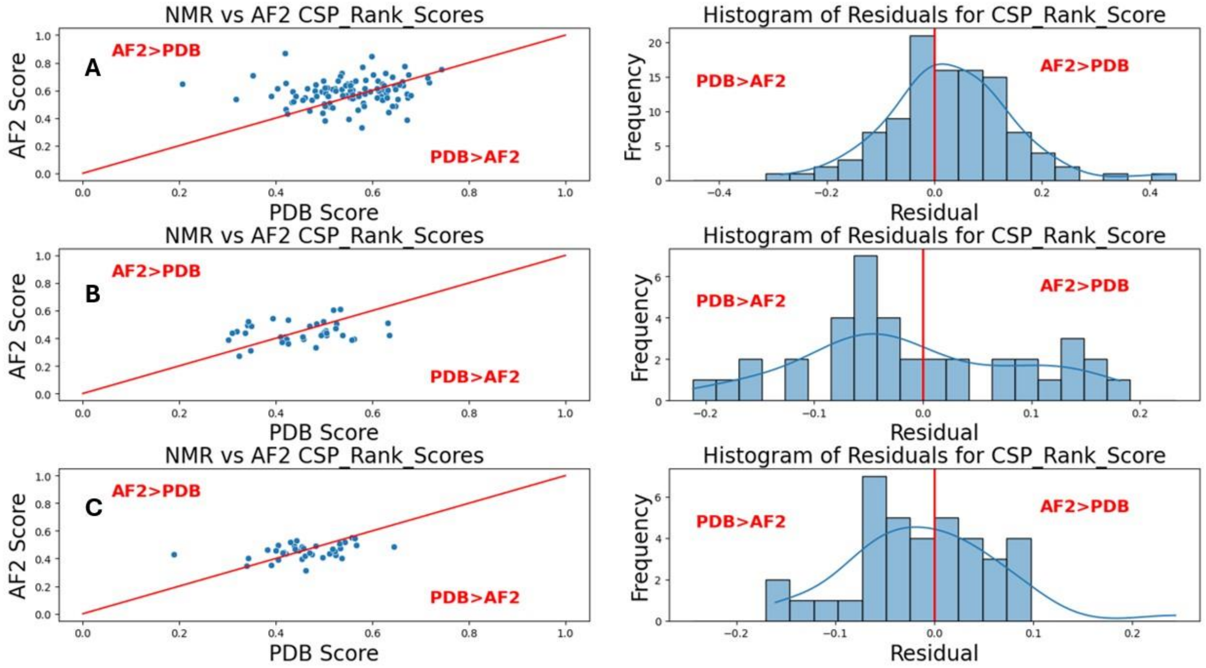


**Fig. S3.** Performance across the dataset using alternative methods of calculating CSPs. A) Method derived from (Williamson, 2013). B) Method derived from (EvenaÈs et al, 2001) C) Method derived from (Grzesiek et al, 1996). These methods of CSP analysis are defined in **Supplementary Note 1**. Due to limitations in data availability, only 38 of the 108 entries have backbone carbon shift data available for the apo and holo forms to generate data for **S2B, C.** This result demonstrates that the distribution of CSP_Rank_Scores changes very little when we use the method derived from (Williamson, 2013). Unfortunately, due to data limitations, it is unclear what relationships can be drawn between the methods which use additional backbone carbon chemical shift data (Grzesiek et al, 1996; EvenaÈs et al, 2001).

**Supplementary Note 1**. Alternative calculations for CSP using carbon chemical shifts in the case of the EVENAES2001 (EvenaÈs et al, 2001) and GRZESIEK1996 (Grzesiek et al, 1996) methods. WILLIAMSON2013 (Williamson, 2013) has different parameters depending on if the residue the shifts are assigned to is Glycine.

WILLIAMSON2013 Method:

If Amino Acid is Glycine: $\text{CSP} = \sqrt{0.5\left((\delta N \cdot 0.20)^2 + \delta H^2\right)}$

Otherwise: $\text{CSP} = \sqrt{0.5\left((\delta N \cdot 0.14)^2 + \delta H^2\right)}$

EVENAES2001 Method: $\text{CSP} = \sqrt{\frac{1}{4}\left(\left(\frac{\delta N}{6.5}\right)^2 + \left(\frac{\delta CA}{3.62}\right)^2 + \left(\frac{\delta CB}{3.62}\right)^2 + \left(\frac{\delta CO}{2.93}\right)^2\right)}$

GRZESIEK1996 Method: $\text{CSP} = \sqrt{\frac{1}{4}\left(\delta H^2 + \left(\frac{\delta N}{5}\right)^2 + \left(\frac{\delta CA}{2}\right)^2 + \left(\frac{\delta CB}{2}\right)^2\right)}$

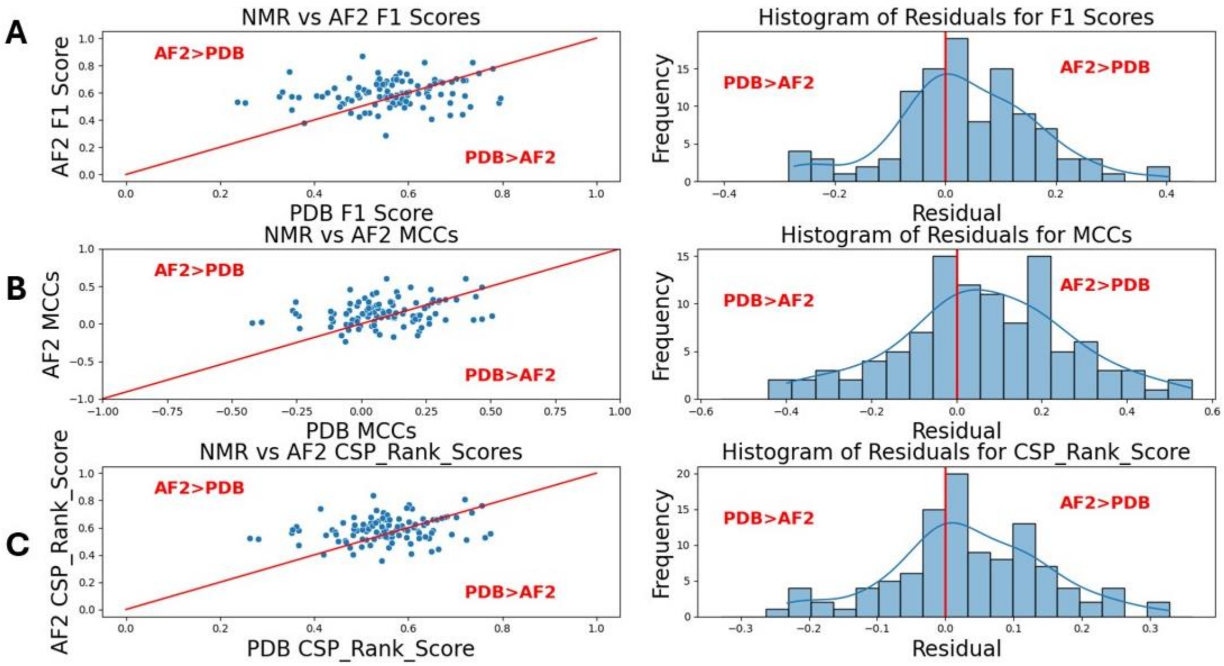


**Fig. S4.** Performance across the dataset for F1, MCC and CSP_Rank_Score statistics. A) Comparison plots for F1 metrics. B) Comparison plots for MCC metrics. C) Comparison plots for CSP_Rank_Score metrics, identical data to **Fig. 1B, C.** CSP_Rank_Score is a weighted average of F1 and MCC scores calculated according to **Equation 6**. For all these metrics, AF2 models (using no sample specific experimental data) perform about as well as (and often better than) models in the PDB determined using CSP and other experimental data to guide the modeling.

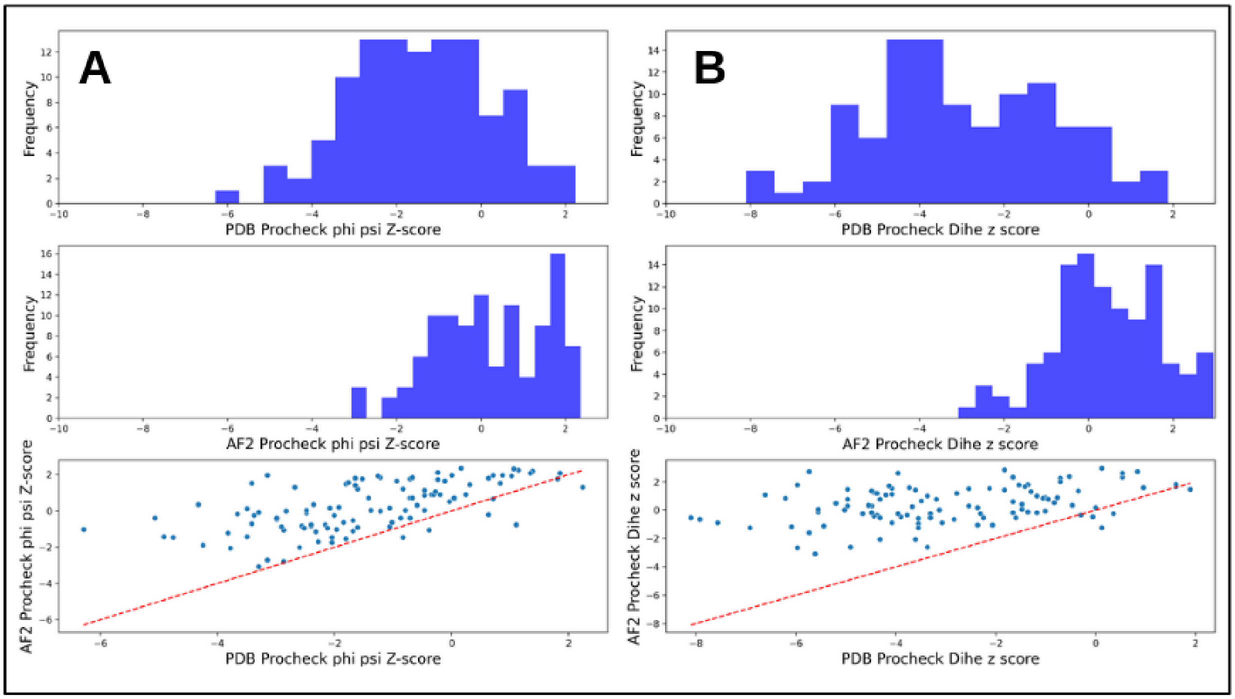


**Fig. S5.** Knowledge based statistical scores between baseline AF2 and medoid NMR models. PSVS v2.0-derived (Bhattacharya et al, 2007) Procheck (Laskowski et al, 1993) comparisons between baseline AF2 models and medoid PDB models. For all of these metrics, AF2 models (using no sample specific experimental data) generally score better than models in the PDB determined using CSP and other experimental data to guide the modeling.

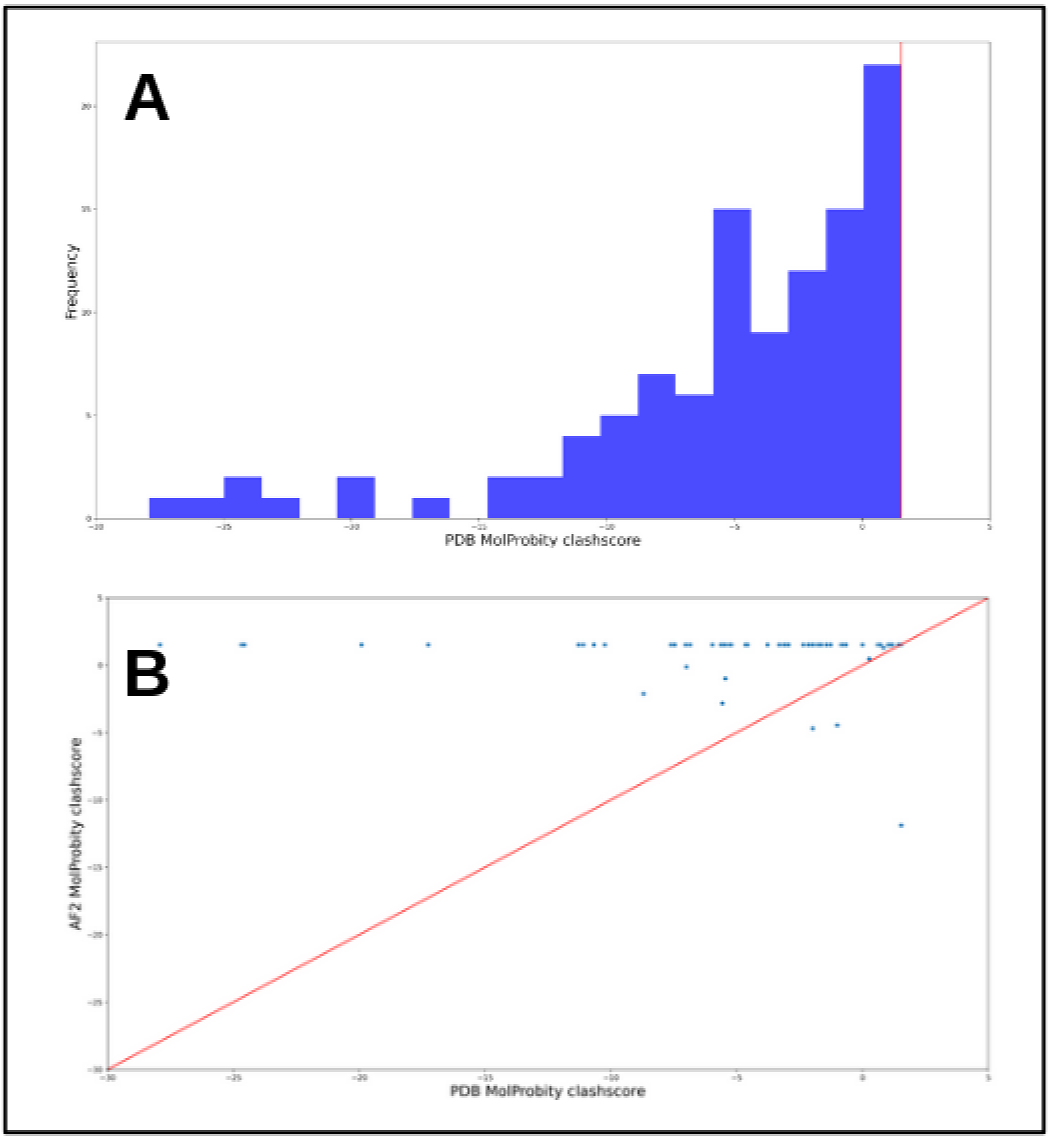


**Fig. S6.** Knowledge based Molprobity clashscores between baseline AF2 and medoid NMR models. PSVS v2.0-derived (Bhattacharya et al, 2007) MolProbity (Chen et al, 2010) clashscore Z-score calculations for medoid holo PDB models. A) Histogram of clashscore Z-scores for medoid PDB models. B) Scatterplot comparing medoid PDB model clashscores Z-scores to those of baseline AF2 models. Of the baseline AF2 models, 98 of the 108 systems have a clashscore of 0 which results in a Z-score of 1.53 (the red vertical line in **S7A)**. The lowest Z-score for a baseline AF2 model was -11.88. In general, AF2 models (using no sample specific experimental data) score significantly better than models in the PDB determined using CSP and other experimental data to guide the modeling.

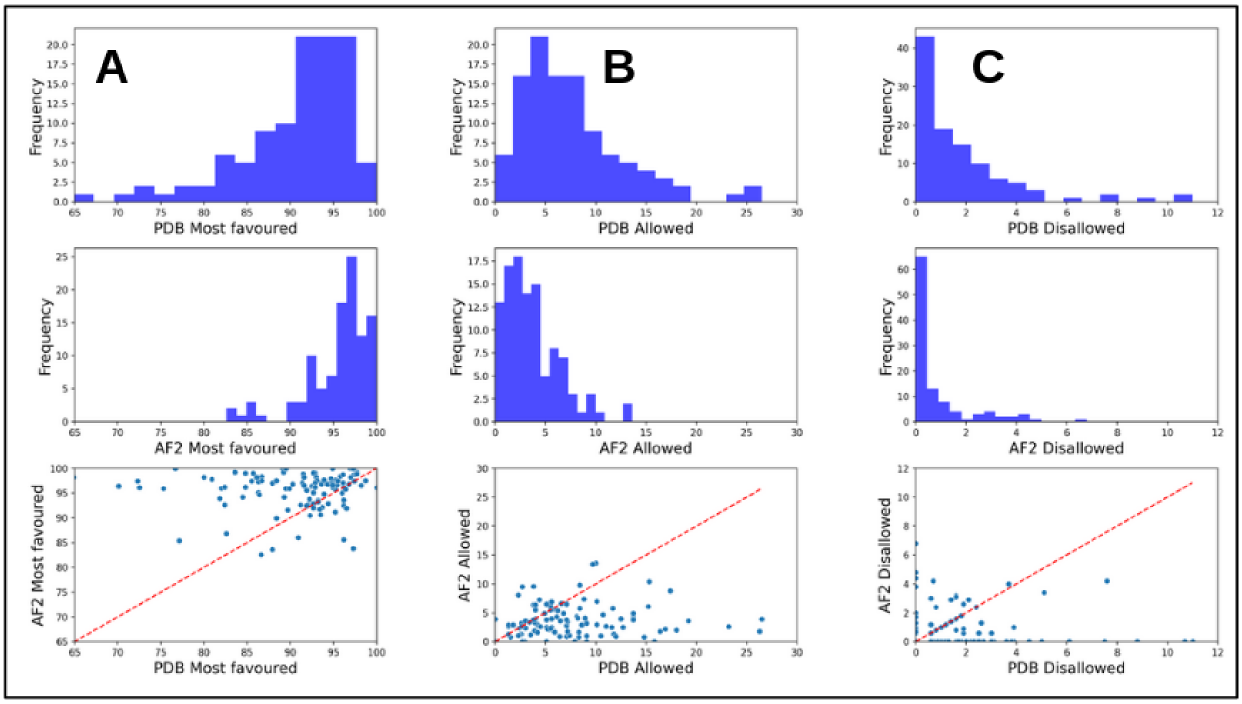


**Fig. S7.** Knowledge based Ramachandran dihedral angle violations between baseline AF2 and medoid NMR models. PSVS v2.0 -derived (Bhattacharya et al, 2007) Ramachandran outlier comparisons (Chen et al, 2010) between baseline AF2 models and medoid PDB models. Overall, AF2 models (using no sample specific experimental data) score significantly better than models in the PDB determined using CSP and other experimental data to guide the modeling, with a shift from phi-psi values in the “allowed” regions of the Ramachandran map to values in the “most favoured” regions.

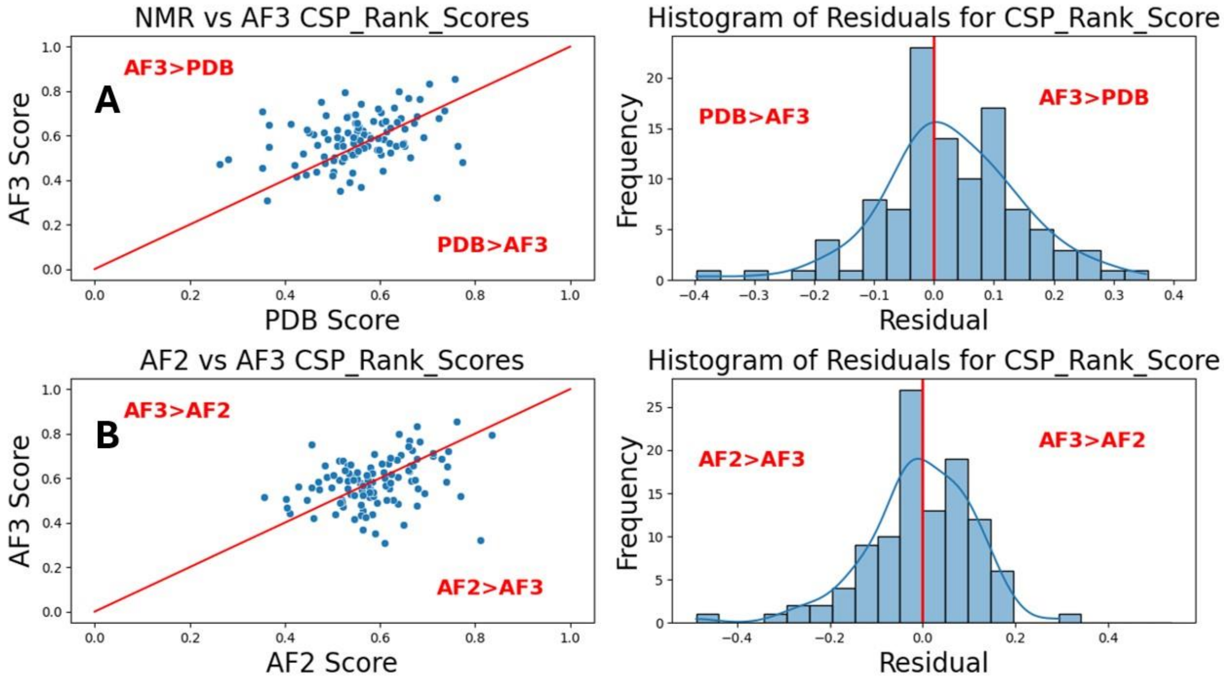


**Fig. S8.** Performance across the dataset using AF3 (Abramson et al, 2024) to generate apo and holo baseline structures as compared to baseline AF2 (Jumper et al, 2021; Evans et al, 2021, Mirdita et al, 2022). A) Comparison between AF3 CSP_Rank_Scores and medoid PDB model scores. B) Comparison between AF3 CSP_Rank_Scores and baseline AF2 scores. This result shows there is little advantage to using baseline AF3 compared to baseline AF2 when assessing the fit of predicted protein-peptide models to CSP data.

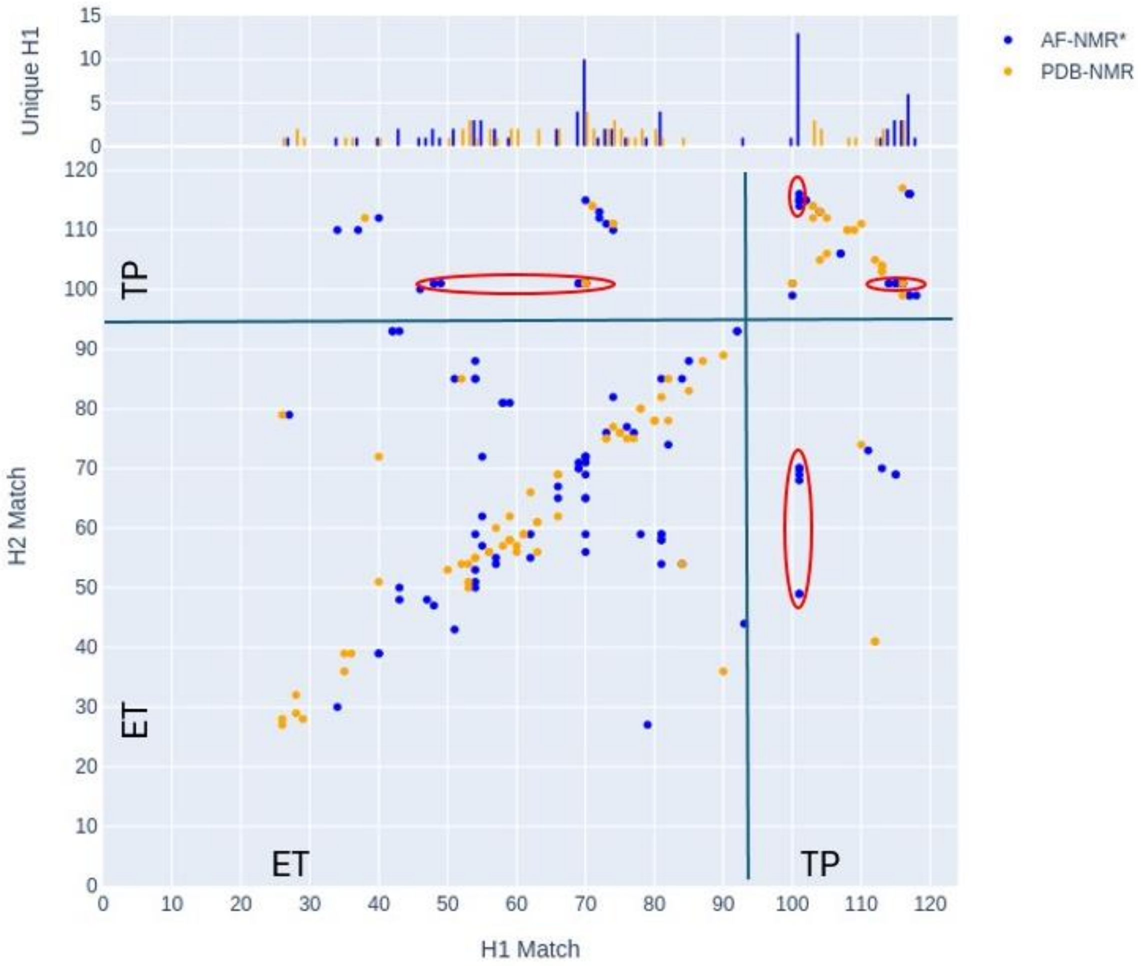


**Fig. S9.** RPF Double Recall analysis for 7JQ8 AF-NMR* and PDB-NMR ensembles. NOEs in the experimental NOESY spectra that can be explained only by the 7JQ8 AF-NMR* ensemble are plotted as blue dots between the residue number of the 1H(-N/C)-donor Match (x-axis) and that of the 1H-acceptor Match (y-axis) (Huang et al, 2021; Huang & Montelione, 2024). NOEs explained only by the 7JQ8 PDB-NMR ensemble are indicated by orange dots. Dots circled in red correspond to NOEs which can be explained by a particular conformation of a TRP (residue 5 of TP) that is consistent with the NMR data for the AF-NMR ensemble but not for the PDB NMR ensemble.

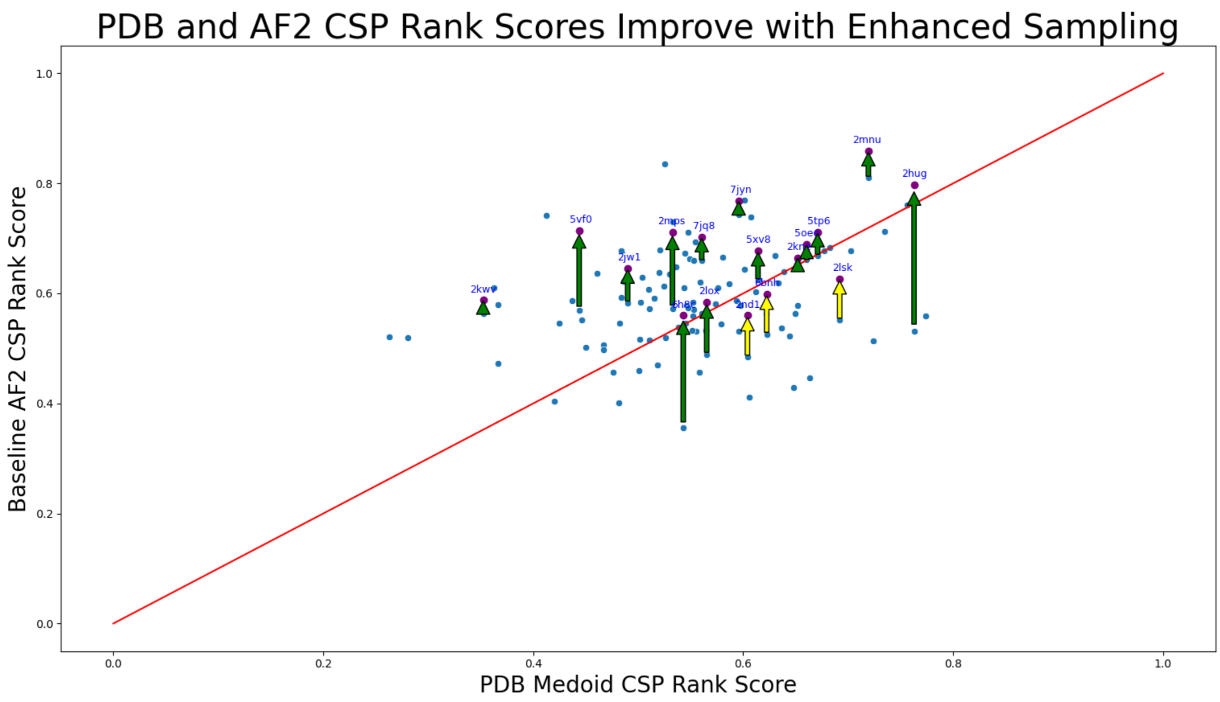


**Fig. S10.** Improvements to baseline AF2 CSP Rank Score using enhanced sampling with AF2 (ES AF2) and the *CSP_Rank* protocol described in the main text. The average improvement in CSP_Rank_Score using the ES AF2 protocol with CSP conformer selection is 0.08 units.

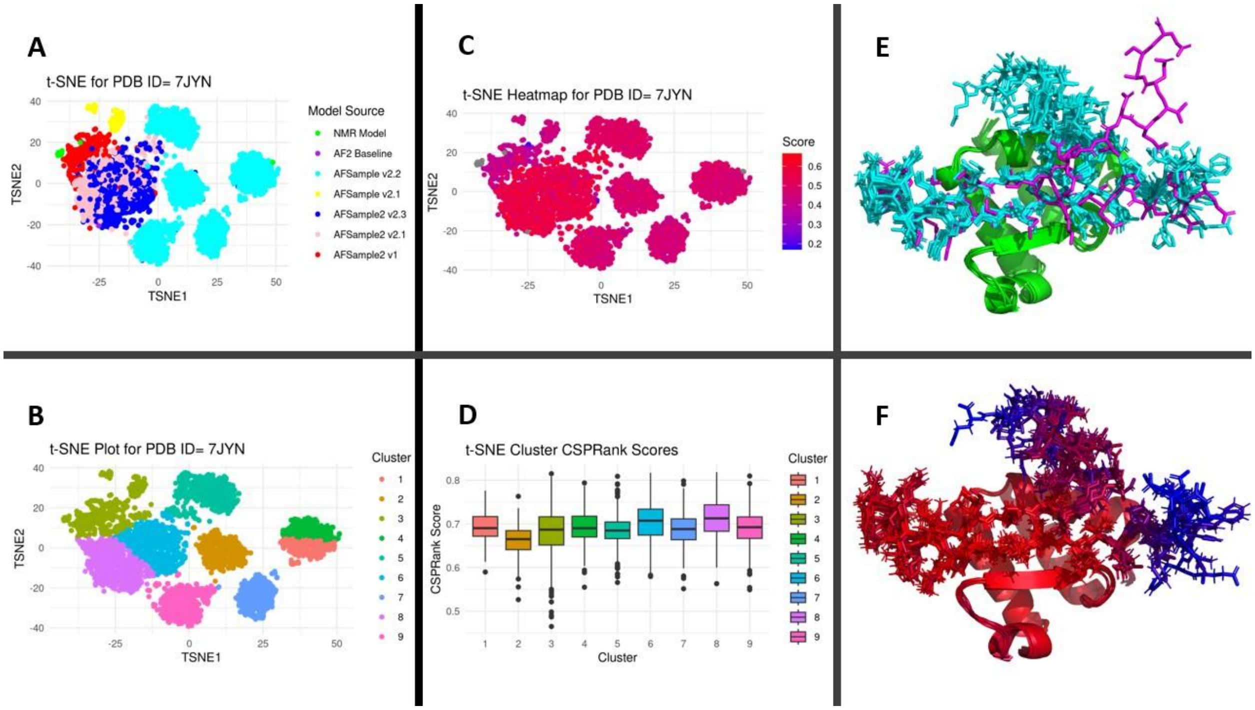


**Fig. S11**. t-SNE clustering analysis and AF-NMR ensemble for 7JYN. Panel legends and coloring are the same as those of the main text **Fig. 4**.

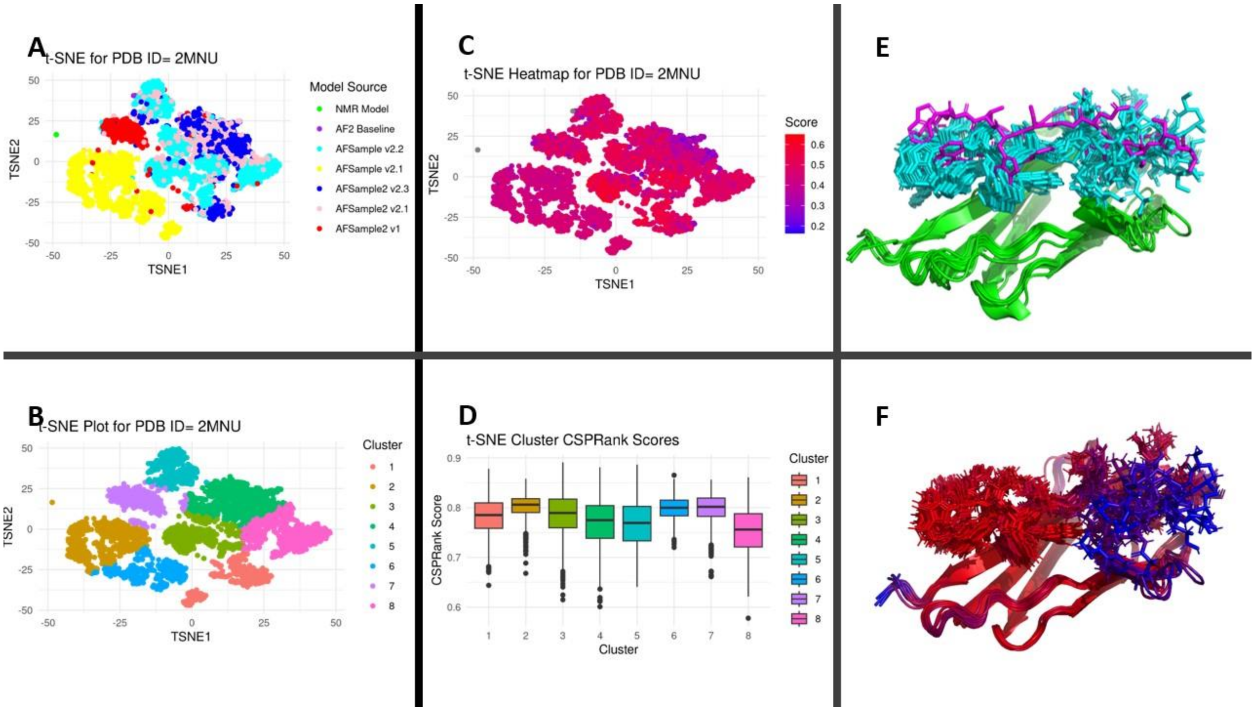


**Fig. S12**. t-SNE clustering analysis and AF-NMR ensemble for 2MNU. Panel legends and coloring are the same as those of the main text **Fig. 4**.

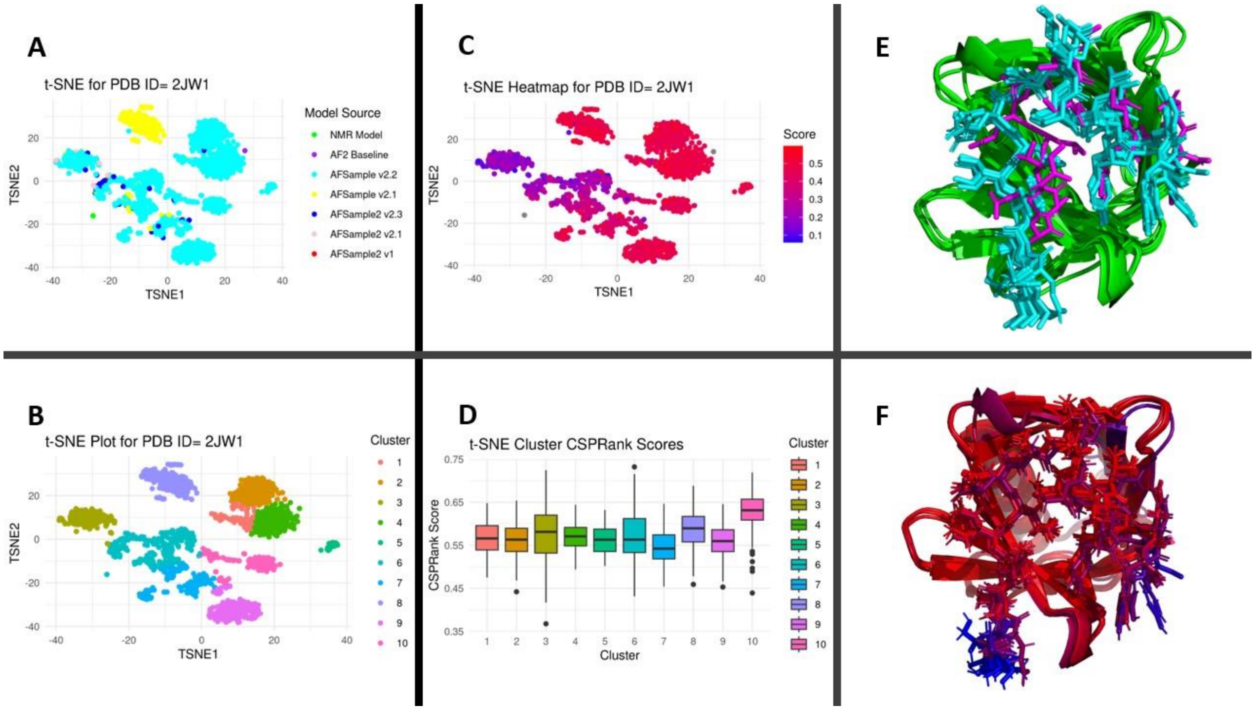


**Fig. S13**. t-SNE clustering analysis and AF-NMR ensemble for 2JW1. Panel legends are the same as those of the main text **Fig. 4**.

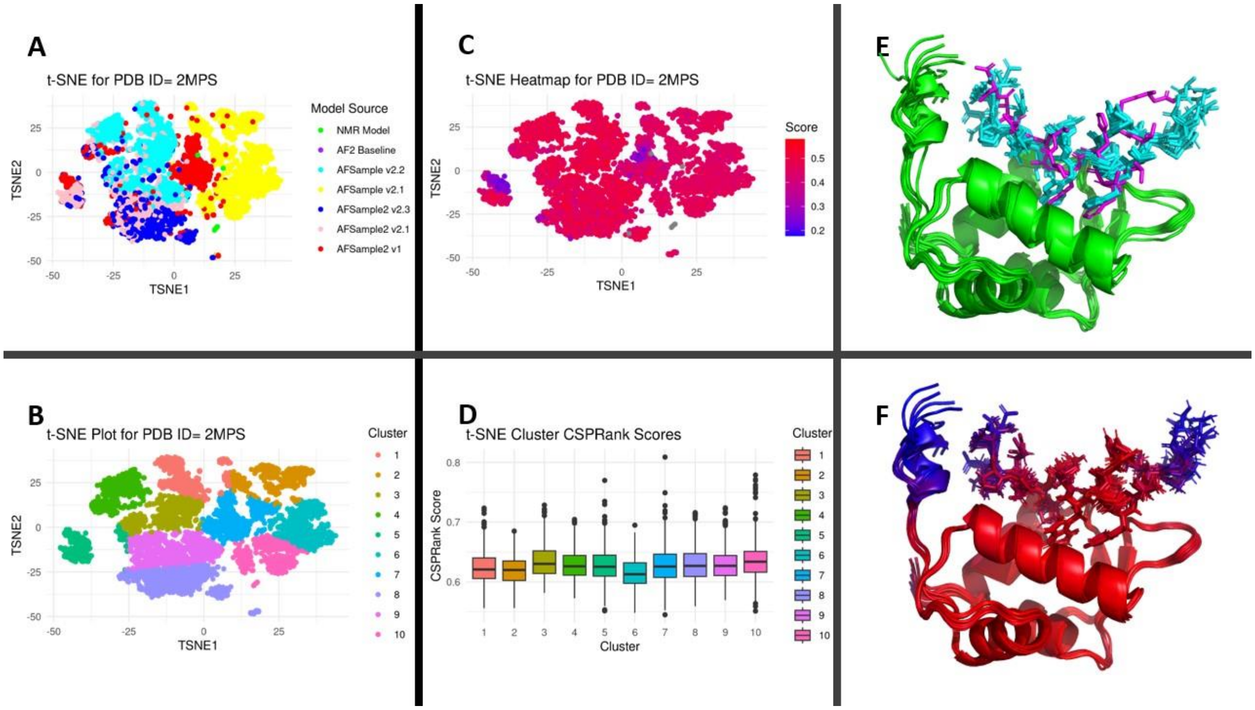


**Fig. S14**. t-SNE clustering analysis and AF-NMR ensemble for 2MPS. Panel legends and coloring are the same as those of the main text **Fig. 4**.

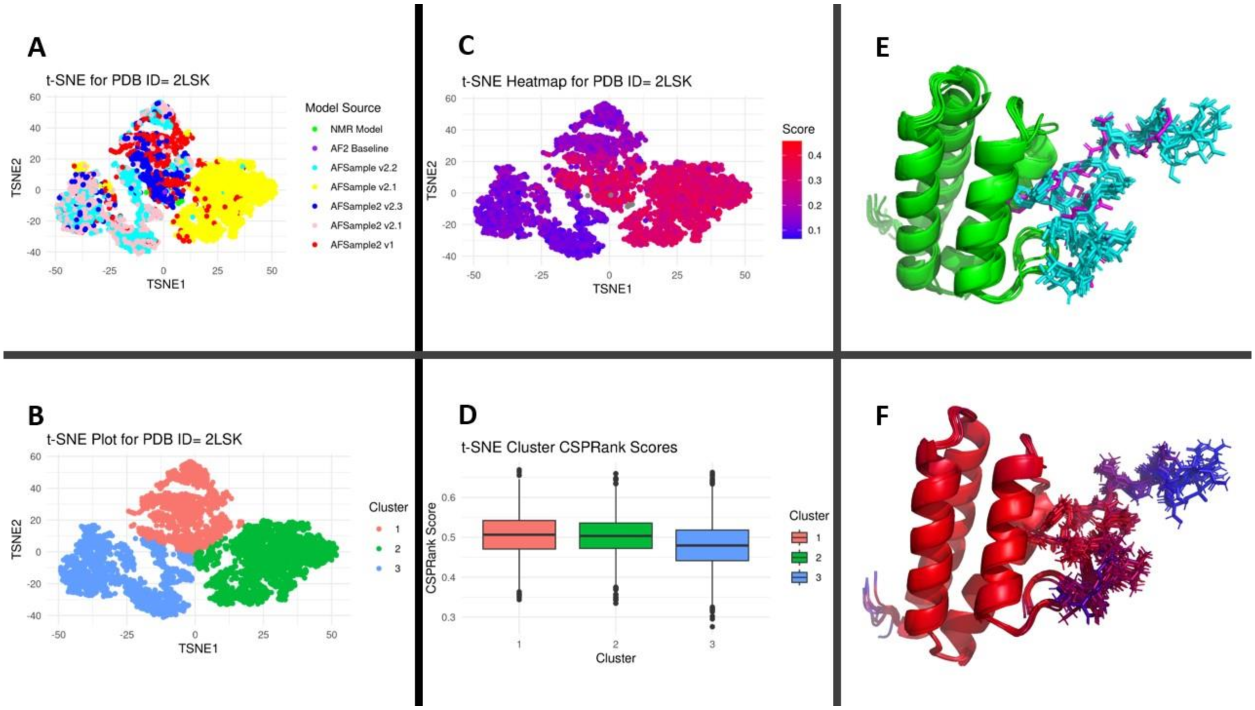


**Fig. S15**. t-SNE clustering analysis and AF-NMR ensemble for 2LSK. Panel legends and coloring are the same as those of the main text **Fig. 4**.

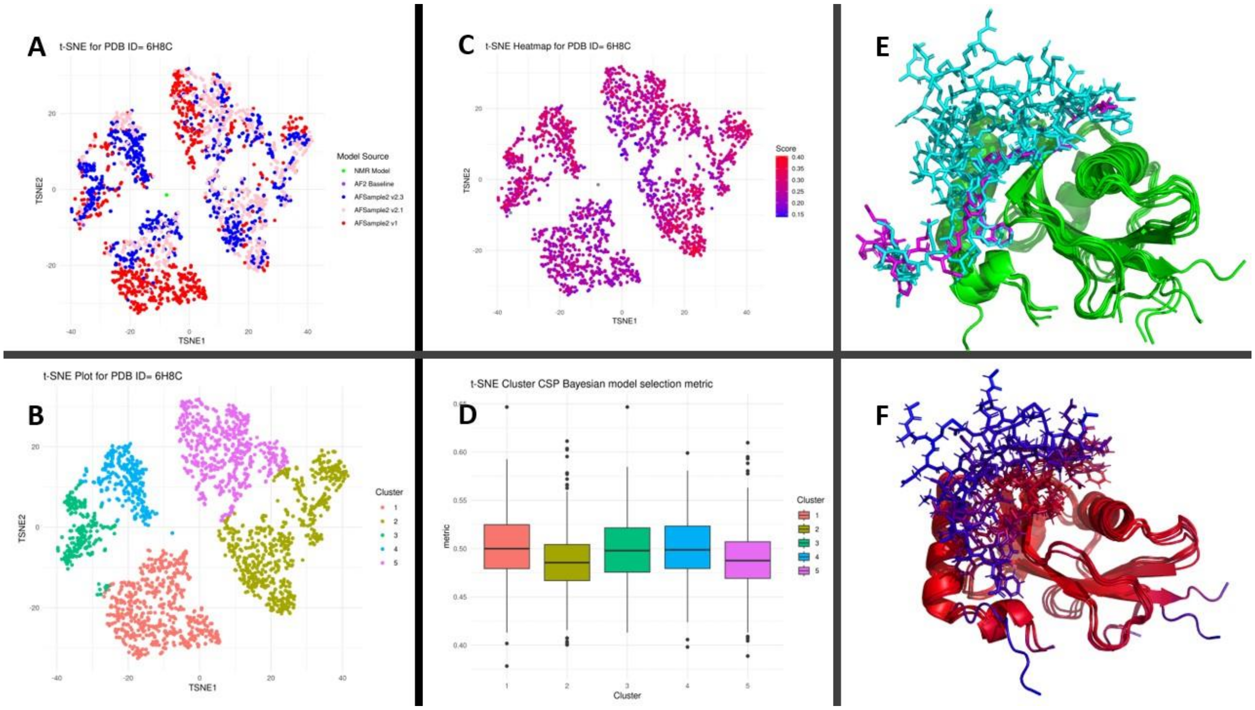


**Fig. S16**. t-SNE clustering analysis and AF-NMR ensemble for 6H8C. Panel legends and coloring are the same as those of the main text **Fig. 4**.

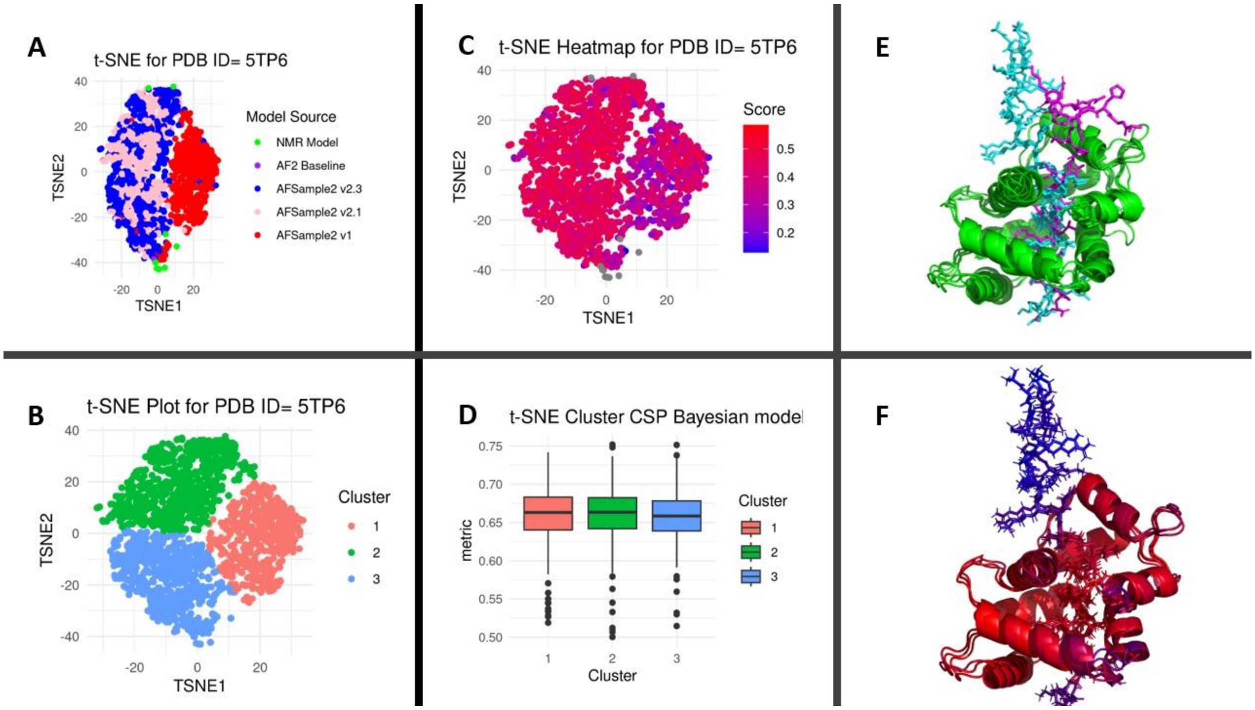


**Fig. S17**. t-SNE clustering analysis and AF-NMR ensemble for 5TP6. Panel legends and coloring are the same as those of the main text **Fig. 4**.

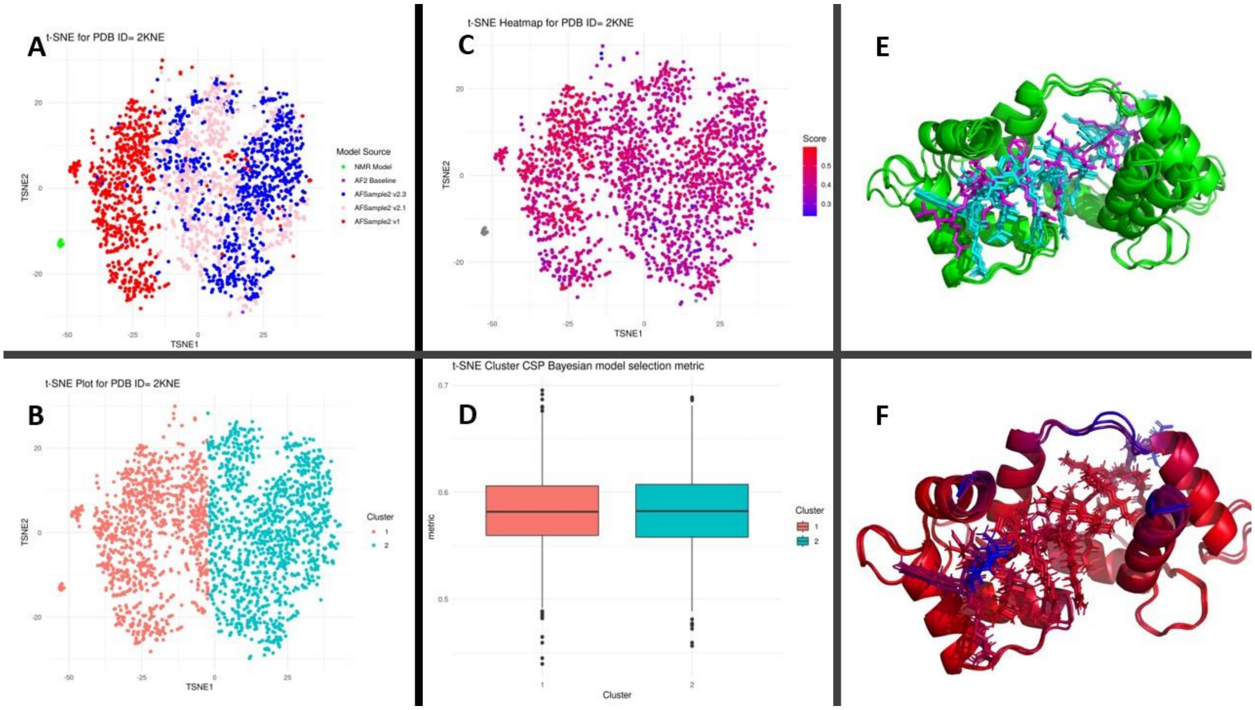


**Fig. S18.** t-SNE clustering analysis and AF-NMR ensemble for 2KNE. Panel legends and coloring are the same as those of the main text **Fig. 4**.

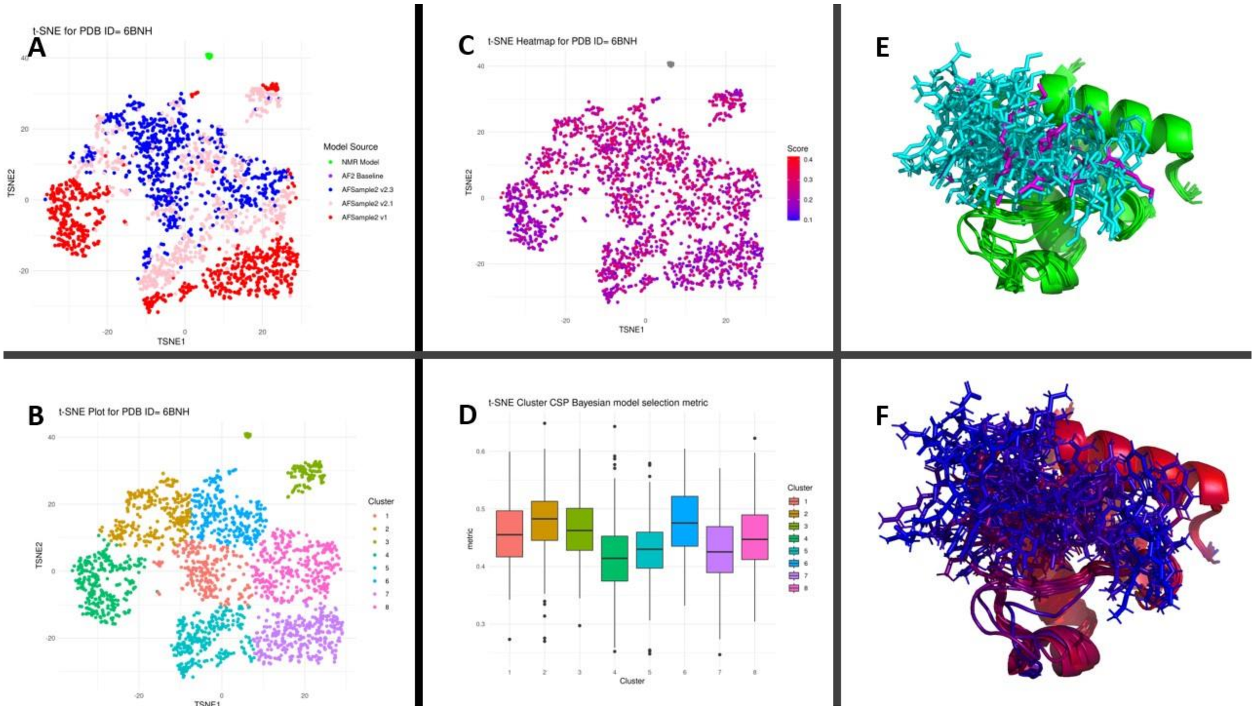


**Fig. S19.** t-SNE clustering analysis and AF-NMR ensemble for 6BNH. Panel legends and coloring are the same as those of the main text **Fig. 4**.

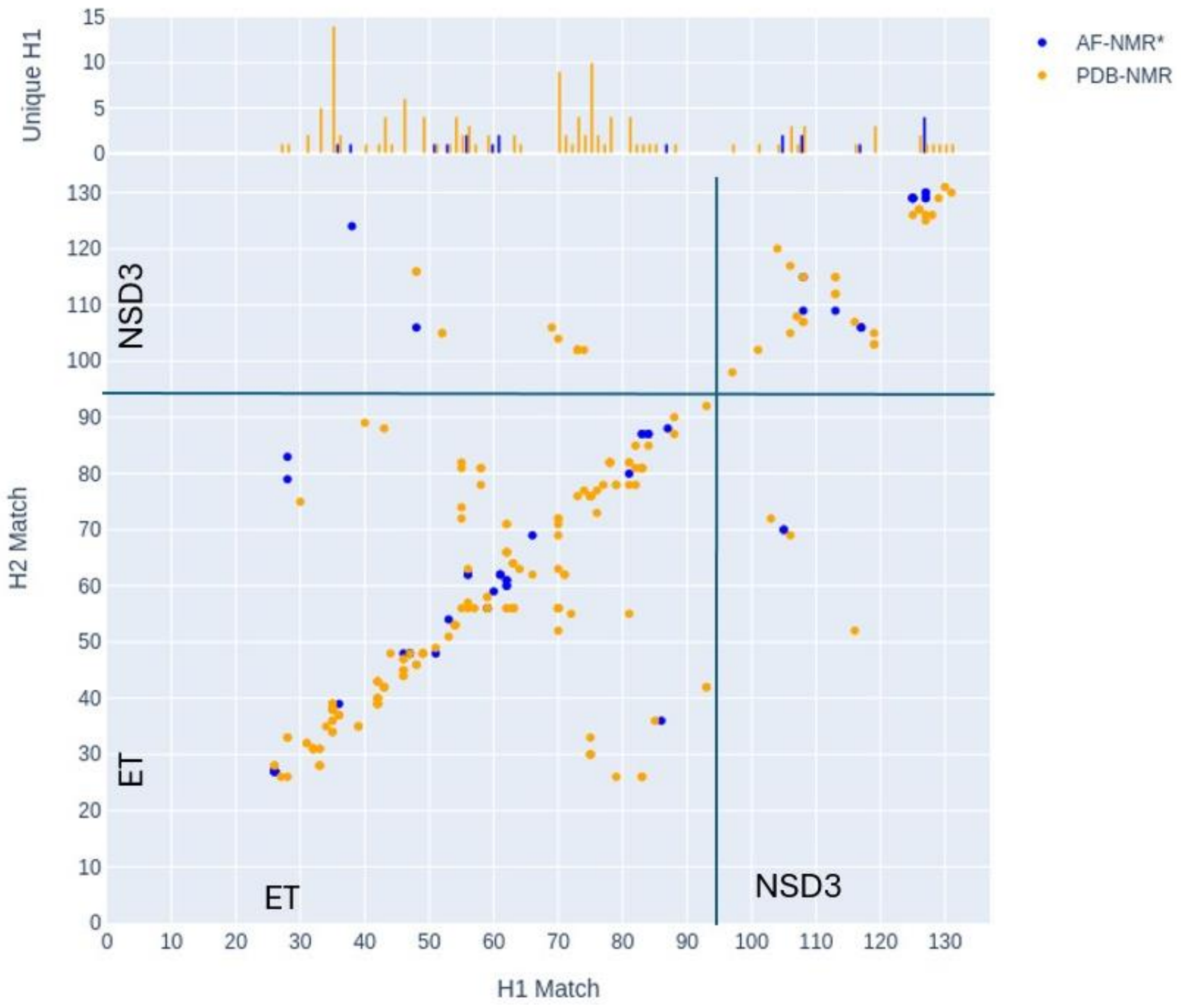


**Fig. S20.** RPF Double Recall analysis for 7JYN AF-NMR* and PDB-NMR ensembles. NOEs in the experimental NOESY spectra that can be explained only by the 7JYN AF-NMR* ensemble are plotted as blue dots between the residue number of the 1H(-N/C)-donor Match (x-axis) and that of the 1H-acceptor Match (y-axis) (Huang et al, 2021; Huang & Montelione, 2024). NOEs explained only by the 7JYN PDB-NMR ensemble are indicated by orange dots.

Analyses performed for DAOP well-defined residues.

| | | | |
|---|---|---|---|
| Total structures computed | currently unknown | | |
| Number of structures used | 11 | | |
| RMSD Values | | | |
| | all | ordered[e] | Selected[f] |
| All backbone atoms | 1.2 Å | 0.5 Å | 0.5 Å |
| All heavy atoms | 1.3 Å | 0.7 Å | 0.7 Å |
| Structure Quality Factors - overall statistics | | | |
| | Mean score | SD | Z-score [g] |
| Procheck G-factor [e] (phi / psi only) | -0.36 | N/A | -1.10 |
| Procheck G-factor [e] (all dihedral angles) | -0.15 | N/A | -0.89 |
| Verify3D | 0.17 | 0.0176 | -4.65 |
| ProsaII (-ve) | 0.56 | 0.0462 | -0.37 |
| MolProbity clashscore | 16.11 | 4.6278 | -1.24 |
| Ramachandran Plot Summary from Procheck [f] | | | |
| Most favoured regions | 94.3% | | |
| Additionally allowed regions | 5.7% | | |
| Generously allowed regions | 0.0% | | |
| Disallowed regions | 0.0% | | |
| Ramachandran Plot Statistics from Richardson's lab | | | |
| Most favoured regions | 97.2% | | |
| Allowed regions | 2.8% | | |
| Disallowed regions | 0% | | |

[e] Residues with sum of phi and psi order parameters >= 1.8: 5M-33M, 36M-70M, 88M-107M

[f] Residues selected (DAOP with S(phi)+S(psi)>=1.8): 5M-33M, 36M-70M, 88M-107M

[g] With respect to mean and standard deviation for a set of 252 X-ray structures < 500 residues, of resolution <= 1.80 Å, R-factor <= 0.25 and R-free <= 0.28; a positive value indicates a 'better' score

Generated using PSVS 2.0-pre

**Supplementary Table S1.** PDB ID 2MNU AF-NMR Ensemble Protein Structure Validation Software Server (Bhattacharya et al, 2007) (PSVS) structure quality factors.

Analyses performed for DAOP well-defined residues.

| | | | |
|---|---|---|---|
| Total structures computed | currently unknown | | |
| Number of structures used | 10 | | |
| RMSD Values | | | |
| | all | ordered[e] | Selected[f] |
| All backbone atoms | 0.4 Å | 0.4 Å | 0.4 Å |
| All heavy atoms | 0.6 Å | 0.6 Å | 0.6 Å |
| Structure Quality Factors - overall statistics | | | |
| | Mean score | SD | Z-score [g] |
| Procheck G-factor [e] (phi / psi only) | -0.06 | N/A | 0.08 |
| Procheck G-factor [e] (all dihedral angles) | 0.13 | N/A | 0.77 |
| Verify3D | 0.18 | 0.0099 | -4.49 |
| ProsaII (-ve) | 0.57 | 0.0577 | -0.33 |
| MolProbity clashscore | 9.50 | 2.6326 | -0.10 |
| Ramachandran Plot Summary from Procheck [f] | | | |
| Most favoured regions | 94.5% | | |
| Additionally allowed regions | 5.5% | | |
| Generously allowed regions | 0.0% | | |
| Disallowed regions | 0.0% | | |
| Ramachandran Plot Statistics from Richardson's lab | | | |
| Most favoured regions | 99.1% | | |
| Allowed regions | 0.9% | | |
| Disallowed regions | 0% | | |

[e] Residues with sum of phi and psi order parameters >= 1.8: 2M-44M, 48M-106M, 118M-133M

[f] Residues selected (DAOP with S(phi)+S(psi)>=1.8): 2M-44M, 48M-106M, 118M-133M

[g] With respect to mean and standard deviation for a set of 252 X-ray structures < 500 residues, of resolution <= 1.80 Å, R-factor <= 0.25 and R-free <= 0.28; a positive value indicates a 'better' score

Generated using PSVS 2.0-pre

**Supplementary Table S2.** PDB ID 2JW1 AF-NMR Ensemble Protein Structure Validation Software Server (Bhattacharya et al, 2007) (PSVS) structure quality factors.

Analyses performed for DAOP well-defined residues.

| | | | |
|---|---|---|---|
| Total structures computed | currently unknown | | |
| Number of structures used | 10 | | |
| RMSD Values | | | |
| | all | ordered[e] | Selected[f] |
| All backbone atoms | 1.0 Å | 0.2 Å | 0.2 Å |
| All heavy atoms | 1.2 Å | 0.5 Å | 0.5 Å |
| Structure Quality Factors - overall statistics | | | |
| | Mean score | SD | Z-score [g] |
| Procheck G-factor [e] (phi / psi only) | 0.16 | N/A | 0.94 |
| Procheck G-factor [e] (all dihedral angles) | 0.24 | N/A | 1.42 |
| Verify3D | 0.21 | 0.0143 | -4.01 |
| ProsaII (-ve) | 0.78 | 0.0395 | 0.54 |
| MolProbity clashscore | 6.48 | 2.2760 | 0.41 |
| Ramachandran Plot Summary from Procheck [f] | | | |
| Most favoured regions | 94.2% | | |
| Additionally allowed regions | 5.8% | | |
| Generously allowed regions | 0.0% | | |
| Disallowed regions | 0.0% | | |
| Ramachandran Plot Statistics from Richardson's lab | | | |
| Most favoured regions | 98.1% | | |
| Allowed regions | 1.9% | | |
| Disallowed regions | 0% | | |

[e] Residues with sum of phi and psi order parameters >= 1.8: 7M-59M, 61M-89M, 104M-112M

[f] Residues selected (DAOP with S(phi)+S(psi)>=1.8): 7M-59M, 61M-89M, 104M-112M

[g] With respect to mean and standard deviation for a set of 252 X-ray structures < 500 residues, of resolution <= 1.80 Å, R-factor <= 0.25 and R-free <= 0.28; a positive value indicates a 'better' score

Generated using PSVS 2.0-pre

**Supplementary Table S3.** PDB ID 2MPS AF-NMR Ensemble Protein Structure Validation Software Server (Bhattacharya et al, 2007) (PSVS) structure quality factors.

Analyses performed for DAOP well-defined residues.

| | | | |
|---|---|---|---|
| Total structures computed | currently unknown | | |
| Number of structures used | 9 | | |
| RMSD Values | | | |
| | all | ordered[e] | Selected[f] |
| All backbone atoms | 0.7 Å | 0.4 Å | 0.4 Å |
| All heavy atoms | 0.8 Å | 0.6 Å | 0.6 Å |
| Structure Quality Factors - overall statistics | | | |
| | Mean score | SD | Z-score [g] |
| Procheck G-factor [e] (phi / psi only) | 0.33 | N/A | 1.61 |
| Procheck G-factor [e] (all dihedral angles) | 0.23 | N/A | 1.36 |
| Verify3D | 0.07 | 0.0120 | -6.26 |
| ProsaII (-ve) | 0.60 | 0.0485 | -0.21 |
| MolProbity clashscore | 12.22 | 3.5484 | -0.57 |
| Ramachandran Plot Summary from Procheck [f] | | | |
| Most favoured regions | 94.3% | | |
| Additionally allowed regions | 5.4% | | |
| Generously allowed regions | 0.2% | | |
| Disallowed regions | 0.0% | | |
| Ramachandran Plot Statistics from Richardson's lab | | | |
| Most favoured regions | 97.6% | | |
| Allowed regions | 1.9% | | |
| Disallowed regions | 0.6% | | |

[e] Residues with sum of phi and psi order parameters >= 1.8: 2M-87M, 102M-115M

[f] Residues selected (DAOP with S(phi)+S(psi)>=1.8): 2M-87M, 102M-115M

[g] With respect to mean and standard deviation for a set of 252 X-ray structures < 500 residues, of resolution <= 1.80 Å, R-factor <= 0.25 and R-free <= 0.28; a positive value indicates a 'better' score

Generated using PSVS 2.0-pre

**Supplementary Table S4.** PDB ID 2LSK AF-NMR Ensemble Protein Structure Validation Software Server (Bhattacharya et al, 2007) (PSVS) structure quality factors.

Analyses performed for DAOP well-defined residues.

| | | | |
|---|---|---|---|
| Total structures computed | currently unknown | | |
| Number of structures used | 6 | | |
| RMSD Values | | | |
| | all | ordered[e] | Selected[f] |
| All backbone atoms | 3.7 Å | 2.0 Å | 2.0 Å |
| All heavy atoms | 4.0 Å | 2.3 Å | 2.3 Å |
| Structure Quality Factors - overall statistics | | | |
| | Mean score | SD | Z-score [g] |
| Procheck G-factor [e] (phi / psi only) | -0.03 | N/A | 0.20 |
| Procheck G-factor [e] (all dihedral angles) | 0.00 | N/A | 0.00 |
| Verify3D | 0.16 | 0.0234 | -4.82 |
| ProsaII (-ve) | 0.79 | 0.0679 | 0.58 |
| MolProbity clashscore | 12.21 | 6.1705 | -0.57 |
| Ramachandran Plot Summary from Procheck [f] | | | |
| Most favoured regions | 92.0% | | |
| Additionally allowed regions | 7.8% | | |
| Generously allowed regions | 0.2% | | |
| Disallowed regions | 0.0% | | |
| Ramachandran Plot Statistics from Richardson's lab | | | |
| Most favoured regions | 97.5% | | |
| Allowed regions | 1.2% | | |
| Disallowed regions | 1.2% | | |

[e] Residues with sum of phi and psi order parameters >= 1.8: 3M-5M, 11M-44M, 48M-110M, 129M-131M, 140M-144M

[f] Residues selected (DAOP with S(phi)+S(psi)>=1.8): 3M-5M, 11M-44M, 48M-110M, 129M-131M, 140M-144M

[g] With respect to mean and standard deviation for a set of 252 X-ray structures < 500 residues, of resolution <= 1.80 Å, R-factor <= 0.25 and R-free <= 0.28; a positive value indicates a 'better' score

Generated using PSVS 2.0-pre

**Supplementary Table S5.** PDB ID 6H8C AF-NMR Ensemble Protein Structure Validation Software Server (Bhattacharya et al, 2007) (PSVS) structure quality factors.

Analyses performed for DAOP well-defined residues.

| | | | |
|---|---|---|---|
| Total structures computed | currently unknown | | |
| Number of structures used | 3 | | |
| RMSD Values | | | |
| | all | ordered[e] | Selected[f] |
| All backbone atoms | 0.8 Å | 0.8 Å | 0.8 Å |
| All heavy atoms | 1.1 Å | 1.0 Å | 1.0 Å |
| Structure Quality Factors - overall statistics | | | |
| | Mean score | SD | Z-score [g] |
| Procheck G-factor [e] (phi / psi only) | 0.31 | N/A | 1.53 |
| Procheck G-factor [e] (all dihedral angles) | 0.38 | N/A | 2.25 |
| Verify3D | 0.15 | 0.0058 | -4.98 |
| ProsaII (-ve) | 1.31 | 0.0300 | 2.73 |
| MolProbity clashscore | 6.29 | 1.8500 | 0.45 |
| Ramachandran Plot Summary from Procheck [f] | | | |
| Most favoured regions | 92.1% | | |
| Additionally allowed regions | 7.9% | | |
| Generously allowed regions | 0.0% | | |
| Disallowed regions | 0.0% | | |
| Ramachandran Plot Statistics from Richardson's lab | | | |
| Most favoured regions | 95.9% | | |
| Allowed regions | 3.7% | | |
| Disallowed regions | 0.4% | | |

[e] Residues with sum of phi and psi order parameters >= 1.8: 2M-73M, 78M-181M

[f] Residues selected (DAOP with S(phi)+S(psi)>=1.8): 2M-73M, 78M-181M

[g] With respect to mean and standard deviation for a set of 252 X-ray structures < 500 residues, of resolution <= 1.80 Å, R-factor <= 0.25 and R-free <= 0.28; a positive value indicates a 'better' score

Generated using PSVS 2.0-pre

**Supplementary Table S6.** PDB ID 5TP6 AF-NMR Ensemble Protein Structure Validation Software Server (Bhattacharya et al, 2007) (PSVS) structure quality factors.

Analyses performed for DAOP well-defined residues.

| | | | |
|---|---|---|---|
| Total structures computed | currently unknown | | |
| Number of structures used | 3 | | |
| RMSD Values | | | |
| | all | ordered[e] | Selected[f] |
| All backbone atoms | 0.4 Å | 0.3 Å | 0.3 Å |
| All heavy atoms | 0.7 Å | 0.6 Å | 0.6 Å |
| Structure Quality Factors - overall statistics | | | |
| | Mean score | SD | Z-score [g] |
| Procheck G-factor [e] (phi / psi only) | 0.44 | N/A | 2.05 |
| Procheck G-factor [e] (all dihedral angles) | 0.47 | N/A | 2.78 |
| Verify3D | 0.18 | 0.0058 | -4.49 |
| ProsaII (-ve) | 1.20 | 0.0100 | 2.27 |
| MolProbity clashscore | 5.02 | 2.5051 | 0.66 |
| Ramachandran Plot Summary from Procheck [f] | | | |
| Most favoured regions | 95.5% | | |
| Additionally allowed regions | 4.5% | | |
| Generously allowed regions | 0.0% | | |
| Disallowed regions | 0.0% | | |
| Ramachandran Plot Statistics from Richardson's lab | | | |
| Most favoured regions | 98% | | |
| Allowed regions | 2% | | |
| Disallowed regions | 0% | | |

[e] Residues with sum of phi and psi order parameters >= 1.8: 2M-70M, 73M-179M

[f] Residues selected (DAOP with S(phi)+S(psi)>=1.8): 2M-70M, 73M-179M

[g] With respect to mean and standard deviation for a set of 252 X-ray structures < 500 residues, of resolution <= 1.80 Å, R-factor <= 0.25 and R-free <= 0.28; a positive value indicates a 'better' score

Generated using PSVS 2.0-pre

**Supplementary Table S7.** PDB ID 2KNE AF-NMR Ensemble Protein Structure Validation Software Server (Bhattacharya et al, 2007) (PSVS) structure quality factors.

Analyses performed for DAOP well-defined residues.

| | | | |
|---|---|---|---|
| Total structures computed | currently unknown | | |
| Number of structures used | 10 | | |
| RMSD Values | | | |
| | all | ordered[e] | Selected[f] |
| All backbone atoms | 1.4 Å | 0.9 Å | 0.9 Å |
| All heavy atoms | 2.0 Å | 1.2 Å | 1.2 Å |
| Structure Quality Factors - overall statistics | | | |
| | Mean score | SD | Z-score [g] |
| Procheck G-factor [e] (phi / psi only) | 0.24 | N/A | 1.26 |
| Procheck G-factor [e] (all dihedral angles) | 0.20 | N/A | 1.18 |
| Verify3D | 0.19 | 0.0278 | -4.33 |
| ProsaII (-ve) | 1.20 | 0.0484 | 2.27 |
| MolProbity clashscore | 25.49 | 15.2217 | -2.85 |
| Ramachandran Plot Summary from Procheck [f] | | | |
| Most favoured regions | 94.2% | | |
| Additionally allowed regions | 5.8% | | |
| Generously allowed regions | 0.0% | | |
| Disallowed regions | 0.0% | | |
| Ramachandran Plot Statistics from Richardson's lab | | | |
| Most favoured regions | 98.4% | | |
| Allowed regions | 1.5% | | |
| Disallowed regions | 0.1% | | |

[e] Residues with sum of phi and psi order parameters >= 1.8: 2M-39M, 41M-43M, 45M-82M

[f] Residues selected (DAOP with S(phi)+S(psi)>=1.8): 2M-39M, 41M-43M, 45M-82M

[g] With respect to mean and standard deviation for a set of 252 X-ray structures < 500 residues, of resolution <= 1.80 Å, R-factor <= 0.25 and R-free <= 0.28; a positive value indicates a 'better' score

Generated using PSVS 2.0-pre

**Supplementary Table S8.** PDB ID 6BNH AF-NMR Ensemble Protein Structure Validation Software Server (Bhattacharya et al, 2007) (PSVS) structure quality factors.

Analyses performed for DAOP well-defined residues.

| | | | |
|---|---|---|---|
| Total structures computed | currently unknown | | |
| Number of structures used | 10 | | |
| RMSD Values | | | |
| | all | ordered[e] | Selected[f] |
| All backbone atoms | 0.4 Å | 0.3 Å | 0.3 Å |
| All heavy atoms | 0.7 Å | 0.6 Å | 0.6 Å |
| Structure Quality Factors - overall statistics | | | |
| | Mean score | SD | Z-score [g] |
| Procheck G-factor [e] (phi / psi only) | 0.21 | N/A | 1.14 |
| Procheck G-factor [e] (all dihedral angles) | 0.35 | N/A | 2.07 |
| Verify3D | 0.21 | 0.0143 | -4.01 |
| MolProbity clashscore | 8.01 | 2.8833 | 0.15 |
| Ramachandran Plot Summary from Procheck [f] | | | |
| Most favoured regions | 97.1% | | |
| Additionally allowed regions | 2.9% | | |
| Generously allowed regions | 0.0% | | |
| Disallowed regions | 0.0% | | |
| Ramachandran Plot Statistics from Richardson's lab | | | |
| Most favoured regions | 100% | | |
| Allowed regions | 0% | | |
| Disallowed regions | 0% | | |

[e] Residues with sum of phi and psi order parameters >= 1.8: 27A-92A, 98A-118A

[f] Residues selected (DAOP with S(phi)+S(psi)>=1.8): 27A-92A, 98A-118A

[g] With respect to mean and standard deviation for a set of 252 X-ray structures < 500 residues, of resolution <= 1.80 Å, R-factor <= 0.25 and R-free <= 0.28; a positive value indicates a 'better' score

Generated using PSVS 2.0-pre

**Supplementary Table S9.** PDB ID 7JQ8 AF-NMR Ensemble Protein Structure Validation Software Server (Bhattacharya et al, 2007) (PSVS) structure quality factors.

Analyses performed for DAOP well-defined residues.

| | | | |
|---|---|---|---|
| Total structures computed | currently unknown | | |
| Number of structures used | 11 | | |
| RMSD Values | | | |
| | all | ordered[e] | Selected[f] |
| All backbone atoms | 0.5 Å | 0.4 Å | 0.4 Å |
| All heavy atoms | 0.8 Å | 0.7 Å | 0.7 Å |
| Structure Quality Factors - overall statistics | | | |
| | Mean score | SD | Z-score [g] |
| Procheck G-factor [e] (phi / psi only) | 0.22 | N/A | 1.18 |
| Procheck G-factor [e] (all dihedral angles) | 0.34 | N/A | 2.01 |
| Verify3D | 0.21 | 0.0130 | -4.01 |
| MolProbity clashscore | 7.05 | 3.3952 | 0.32 |
| Ramachandran Plot Summary from Procheck [f] | | | |
| Most favoured regions | 97.6% | | |
| Additionally allowed regions | 2.4% | | |
| Generously allowed regions | 0.0% | | |
| Disallowed regions | 0.0% | | |
| Ramachandran Plot Statistics from Richardson's lab | | | |
| Most favoured regions | 100% | | |
| Allowed regions | 0% | | |
| Disallowed regions | 0% | | |

[e] Residues with sum of phi and psi order parameters >= 1.8: 27A-92A, 98A-118A

[f] Residues selected (DAOP with S(phi)+S(psi)>=1.8): 27A-92A, 98A-118A

[g] With respect to mean and standard deviation for a set of 252 X-ray structures < 500 residues, of resolution <= 1.80 Å, R-factor <= 0.25 and R-free <= 0.28; a positive value indicates a 'better' score

Generated using PSVS 2.0-pre

**Supplementary Table S10.** PDB ID 7JQ8 AF-NMR* (joint fitting to CSP and NOESY data) Ensemble Protein Structure Validation Software Server (Bhattacharya et al, 2007) (PSVS) structure quality factors.

Analyses performed for DAOP well-defined residues.

| | | | |
|---|---|---|---|
| Total structures computed | currently unknown | | |
| Number of structures used | 20 | | |
| RMSD Values | | | |
| | all | ordered[e] | Selected[f] |
| All backbone atoms | 6.9 Å | 0.6 Å | 0.6 Å |
| All heavy atoms | 7.4 Å | 1.1 Å | 1.1 Å |
| Structure Quality Factors - overall statistics | | | |
| | Mean score | SD | Z-score [g] |
| Procheck G-factor [e] (phi / psi only) | -0.03 | N/A | 0.20 |
| Procheck G-factor [e] (all dihedral angles) | -0.13 | N/A | -0.77 |
| Verify3D | 0.13 | 0.0105 | -5.30 |
| ProsaII (-ve) | 0.57 | 0.0552 | -0.33 |
| MolProbity clashscore | 6.01 | 2.1394 | 0.49 |
| Ramachandran Plot Summary from Procheck [f] | | | |
| Most favoured regions | 92.1% | | |
| Additionally allowed regions | 7.9% | | |
| Generously allowed regions | 0.0% | | |
| Disallowed regions | 0.0% | | |
| Ramachandran Plot Statistics from Richardson's lab | | | |
| Most favoured regions | 97.8% | | |
| Allowed regions | 2.2% | | |
| Disallowed regions | 0% | | |

[e] Residues with sum of phi and psi order parameters >= 1.8: 26M-93M, 112M-115M, 118M-126M

[f] Residues selected (DAOP with S(phi)+S(psi)>=1.8): 26M-93M, 112M-115M, 118M-126M

[g] With respect to mean and standard deviation for a set of 252 X-ray structures < 500 residues, of resolution <= 1.80 Å, R-factor <= 0.25 and R-free <= 0.28; a positive value indicates a 'better' score

Generated using PSVS 2.0-pre

**Supplementary Table S11.** PDB ID 7JQ8 NMR Ensemble (directly from PDB) Protein Structure Validation Software Server (Bhattacharya et al, 2007) (PSVS) structure quality factors.

Analyses performed for DAOP well-defined residues.

| | | | |
|---|---|---|---|
| Total structures computed | currently unknown | | |
| Number of structures used | 8 | | |
| RMSD Values | | | |
| | all | ordered[e] | Selected[f] |
| All backbone atoms | 0.8 Å | 0.7 Å | 0.7 Å |
| All heavy atoms | 1.1 Å | 1.0 Å | 1.0 Å |
| Structure Quality Factors - overall statistics | | | |
| | Mean score | SD | Z-score [g] |
| Procheck G-factor [e] (phi / psi only) | 0.02 | N/A | 0.39 |
| Procheck G-factor [e] (all dihedral angles) | 0.09 | N/A | 0.53 |
| Verify3D | 0.20 | 0.0151 | -4.17 |
| ProsaII (-ve) | 0.86 | 0.0389 | 0.87 |
| MolProbity clashscore | 13.35 | 4.4343 | -0.77 |
| Ramachandran Plot Summary from Procheck [f] | | | |
| Most favoured regions | 93.5% | | |
| Additionally allowed regions | 6.5% | | |
| Generously allowed regions | 0.0% | | |
| Disallowed regions | 0.0% | | |
| Ramachandran Plot Statistics from Richardson's lab | | | |
| Most favoured regions | 97% | | |
| Allowed regions | 2.6% | | |
| Disallowed regions | 0.4% | | |

[e] Residues with sum of phi and psi order parameters >= 1.8: 27A-78A, 81A-93A, 95A-131A

[f] Residues selected (DAOP with S(phi)+S(psi)>=1.8): 27A-78A, 81A-93A, 95A-131A

[g] With respect to mean and standard deviation for a set of 252 X-ray structures < 500 residues, of resolution <= 1.80 Å, R-factor <= 0.25 and R-free <= 0.28; a positive value indicates a 'better' score

Generated using PSVS 2.0-pre

**Supplementary Table S12.** PDB ID 7JYN AF-NMR Ensemble Protein Structure Validation Software Server (Bhattacharya et al, 2007) (PSVS) structure quality factors.

Analyses performed for DAOP well-defined residues.

| | | | |
|---|---|---|---|
| Total structures computed | currently unknown | | |
| Number of structures used | 8 | | |
| RMSD Values | | | |
| | all | ordered[e] | Selected[f] |
| All backbone atoms | 0.9 Å | 0.8 Å | 0.8 Å |
| All heavy atoms | 1.2 Å | 1.1 Å | 1.1 Å |
| Structure Quality Factors - overall statistics | | | |
| | Mean score | SD | Z-score [g] |
| Procheck G-factor [e] (phi / psi only) | 0.05 | N/A | 0.51 |
| Procheck G-factor [e] (all dihedral angles) | 0.10 | N/A | 0.59 |
| Verify3D | 0.20 | 0.0173 | -4.17 |
| ProsaII (-ve) | 0.88 | 0.0327 | 0.95 |
| MolProbity clashscore | 13.42 | 2.4461 | -0.78 |
| Ramachandran Plot Summary from Procheck [f] | | | |
| Most favoured regions | 93.5% | | |
| Additionally allowed regions | 6.5% | | |
| Generously allowed regions | 0.0% | | |
| Disallowed regions | 0.0% | | |
| Ramachandran Plot Statistics from Richardson's lab | | | |
| Most favoured regions | 97.2% | | |
| Allowed regions | 2.7% | | |
| Disallowed regions | 0.1% | | |

[e] Residues with sum of phi and psi order parameters >= 1.8: 27A-93A, 95A-131A

[f] Residues selected (DAOP with S(phi)+S(psi)>=1.8): 27A-93A, 95A-131A

[g] With respect to mean and standard deviation for a set of 252 X-ray structures < 500 residues, of resolution <= 1.80 Å, R-factor <= 0.25 and R-free <= 0.28; a positive value indicates a 'better' score

Generated using PSVS 2.0-pre

**Supplementary Table S13.** 7JYN AF-NMR* Ensemble (joint fitting to CSP and NOESY data) Protein Structure Validation Software Server (Bhattacharya et al, 2007) (PSVS) structure quality factors.

Analyses performed for DAOP well-defined residues.

| | | | |
|---|---|---|---|
| Total structures computed | currently unknown | | |
| Number of structures used | 20 | | |
| RMSD Values | | | |
| | all | ordered[e] | Selected[f] |
| All backbone atoms | 8.5 Å | 0.6 Å | 0.6 Å |
| All heavy atoms | 8.8 Å | 1.1 Å | 1.1 Å |
| Structure Quality Factors - overall statistics | | | |
| | Mean score | SD | Z-score [g] |
| Procheck G-factor [e] (phi / psi only) | -0.24 | N/A | -0.63 |
| Procheck G-factor [e] (all dihedral angles) | -0.24 | N/A | -1.42 |
| Verify3D | 0.10 | 0.0140 | -5.78 |
| ProsaII (-ve) | 0.42 | 0.0609 | -0.95 |
| MolProbity clashscore | 6.36 | 2.0756 | 0.43 |
| Ramachandran Plot Summary from Procheck [f] | | | |
| Most favoured regions | 84.5% | | |
| Additionally allowed regions | 15.5% | | |
| Generously allowed regions | 0.0% | | |
| Disallowed regions | 0.0% | | |
| Ramachandran Plot Statistics from Richardson's lab | | | |
| Most favoured regions | 95.3% | | |
| Allowed regions | 4.6% | | |
| Disallowed regions | 0.1% | | |

[e] Residues with sum of phi and psi order parameters >= 1.8: 26M-93M, 114M-120M, 127M-132M

[f] Residues selected (DAOP with S(phi)+S(psi)>=1.8): 26M-93M, 114M-120M, 127M-132M

[g] With respect to mean and standard deviation for a set of 252 X-ray structures < 500 residues, of resolution <= 1.80 Å, R-factor <= 0.25 and R-free <= 0.28; a positive value indicates a 'better' score

Generated using PSVS 2.0-pre

**Supplementary Table S14.** PDB ID 7JYN NMR Ensemble (directly from PDB) Protein Structure Validation Software Server (Bhattacharya et al, 2007) (PSVS) structure quality factors.

### Supplementary Information References


Abramson, J., Adler, J., Dunger, J., Evans, R., Green, T., Pritzel, A., Ronneberger, O., Willmore, L., Ballard, A.J., Bambrick, J. and Bodenstein, S.W., 2024. Accurate structure prediction of biomolecular interactions with AlphaFold 3. *Nature*, pp.1-3. https://doi.org/10.1038/s41586-024-07487-w

Basu, S. and Wallner, B., 2016. DockQ: a quality measure for protein-protein docking models. *PLOS One*, *11*(8), p.e0161879. https://doi.org/10.1371/journal.pone.0161879

Bhattacharya, A., Tejero, R. and Montelione, G.T., 2007. Evaluating protein structures determined by structural genomics consortia. *PROTEINS: Struct Funct Bioinformatics*, *66*(4), pp.778-795. https://doi.org/10.1002/prot.21165

Chen, V.B., Arendall, W.B., Headd, J.J., Keedy, D.A., Immormino, R.M., Kapral, G.J., Murray, L.W., Richardson, J.S. and Richardson, D.C., 2010. MolProbity: all-atom structure validation for macromolecular crystallography. Acta Crystall Section D: Biological Crystallography, *66*(1), pp.12-21.https://doi.org/10.1107/97809553602060000884

Evans, R., O'Neill, M., Pritzel, A., Antropova, N., Senior, A., Green, T., Žídek, A., Bates, R., Blackwell, S., Yim, J. and Ronneberger, O., 2021. Protein complex prediction with AlphaFold-Multimer. *biorxiv*, pp.2021-10. https://doi.org/10.1101/2021.10.04.463034

EvenaÈs, J., Tugarinov, V., Skrynnikov, N.R., Goto, N.K., Muhandiram, R. and Kay, L.E., 2001. Ligand-induced structural changes to maltodextrin-binding protein as studied by solution NMR spectroscopy. *J Mol Biol*, *309*(4), pp.961-974. https://doi.org/10.1006/jmbi.2001.4695

Grzesiek, S., Bax, A., Clore, G.M., Gronenborn, A.M., Hu, J.S., Kaufman, J., Palmer, I., Stahl, S.J. and Wingfield, P.T., 1996. The solution structure of HIV-1 Nef reveals an unexpected fold and permits delineation of the binding surface for the SH3 domain of Hck tyrosine protein kinase. *Nat Struct Biol*, *3*(4), pp.340-345. https://doi.org/10.1038/nsb0496-340

Han, B., Liu, Y., Ginzinger, S.W. and Wishart, D.S., 2011. SHIFTX2: significantly improved protein chemical shift prediction. *J. Biomol. NMR*, *50*, pp.43-57. https://doi.org/10.1007/s10858-011-9478-4

Huang, Y.J., Zhang, N., Bersch, B., Fidelis, K., Inouye, M., Ishida, Y., Kryshtafovych, A., Kobayashi, N., Kuroda, Y., Liu, G. and LiWang, A., 2021. Assessment of prediction methods for protein structures determined by NMR in CASP14: Impact of AlphaFold2. *PROTEINS: Struct Funct Bioinformatics*, *89*(12), pp.1959-1976. https://doi.org/10.1002/prot.26246

Huang, Y.J. and Montelione, G.T., 2024. Hidden Structural States of Proteins Revealed by Conformer Selection with AlphaFold-NMR. *BioRxiv*. https://doi.org/10.1101/2024.06.26.600902

Kirchner, D.K. and Güntert, P., 2011. Objective identification of residue ranges for the superposition of protein structures. *BMC bioinformatics*, *12*, pp.1-11. https://doi.org/10.1186/1471-2105-12-170

Laskowski, R.A., MacArthur, M.W., Moss, D.S. and Thornton, J.M., 1993. PROCHECK: a program to check the stereochemical quality of protein structures. *Journal of applied crystallography*, *26*(2), pp.283-291. https://doi.org/10.1107/s0021889892009944

Li, J., Bennett, K.C., Liu, Y., Martin, M.V. and Head-Gordon, T., 2020. Accurate prediction of chemical shifts for aqueous protein structure on "Real World" data. *Chem. Sci. 11*(12), pp.3180-3191. https://doi.org/10.1039/c9sc06561j

Jumper, J., Evans, R., Pritzel, A., Green, T., Figurnov, M., Ronneberger, O., Tunyasuvunakool, K., Bates, R., Žídek, A., Potapenko, A. and Bridgland, A., 2021. Highly accurate protein structure prediction with AlphaFold. *Nature*, *596*(7873), pp.583-589. https://doi.org/10.1038/s41586-021-03819-2

Mirdita, M., Schütze, K., Moriwaki, Y., Heo, L., Ovchinnikov, S. and Steinegger, M., 2022. ColabFold: making protein folding accessible to all. *Nature methods*, *19*(6), pp.679-682.

https://doi.org/10.1038/s41592-022-01488-1
Shen, Y. and Bax, A., 2010. SPARTA+: a modest improvement in empirical NMR chemical shift prediction by means of an artificial neural network. *J Biomol. NMR*, *48*, pp.13-22. https://doi.org/10.1007/s10858-010-9433-9
Snyder, D.A., Grullon, J., Huang, Y.J., Tejero, R. and Montelione, G.T., 2014. The expanded FindCore method for identification of a core atom set for assessment of protein structure prediction. *PROTEINS: Struct Funct Bioinformatics*, *82*, pp.219-230. https://doi.org/10.1002/prot.24490
Williamson, M.P., 2013. Using chemical shift perturbation to characterise ligand binding. *Prog. Nucl. Magn. Reson. Spectrosc.* *73*, pp.1-16. https://doi.org/10.1016/j.pnmrs.2014.05.001
Zhang, Y. and Skolnick, J., 2005. TM-align: a protein structure alignment algorithm based on the TM-score. *Nucl Acids Res*, *33*(7), pp.2302-2309. https://doi.org/10.1093/nar/gki524
